# Lariat debranching by RNA DEBRANCHING ENZYME 1 depends on SICKLE in *Arabidopsis thaliana*

**DOI:** 10.1101/2023.07.25.550414

**Authors:** Emma E. Kovak, Carine M. Marshall, Mayla D. C. Molinari, Alexandre L. Nepomuceno, Frank G. Harmon

## Abstract

Spliceosome mediated intron removal from precursor mRNAs (pre-mRNAs) generates circular RNAs called intron lariats. RNA DEBRANCHING ENZYME 1 (DBR1) ribonucleases linearize, or debranch, intron lariats to allow their degradation. *DBR1* genes occur across eukaryotes and are essential in animals and plants. High levels of intron lariats in the weak *Arabidopsis thaliana dbr1-2* allele inhibits primary microRNA (pri-miRNA) processing, disrupting miRNA production and miRNA-regulated growth and development. Arabidopsis *sickle* (*sic*) mutants alter pri-miRNA processing and pre-mRNA splicing. This study demonstrates *sic* mutants accumulate intron lariats matching those in weak *dbr* alleles. The strong *sic-1* and weak *dbr1-3* alleles together cause synthetic lethality, while weak *sic-3* with *dbr1-3* has intron lariat accumulation like *sic-3*. Further, *sic-3*, *dbr1-3*, and *sic-3 dbr1-3* similarly alter circadian rhythms and growth of roots and rosettes. The conserved MPLKIP amino acid motif in SIC mediates physical interaction with DBR1 *in vitro* and is required for intron lariat debranching *in vivo*. Thus, MPLKIP containing proteins, like SIC and human TTDN1, act with cognate DBR1 proteins to maintain RNA homeostasis critical for growth and development.

## INTRODUCTION

The spliceosome is the multisubunit ribonucleoprotein complex that executes splicing of precursor messenger RNAs (pre-mRNAs) to generate mRNAs. The spliceosome assembles *de novo* at each exon-intron-exon region of a pre-mRNA to remove the intron and join the flanking exons (Shi, 2017). Intron lariats are circular RNAs derived from introns that are a conserved byproduct of the splicing reaction (Grabowski et al., 1984; Padgett et al., 1984; Ruskin et al., 1984). The intron lariat structure consists of a circular region, which is formed by an unusual 2’-5’ phosphodiester bond between the 5’-phosphate at the 5’-end of the intron and the 2’-hydoxyl of an internal intron nucleotide at a site called the branchpoint, and a linear single-stranded tail with a free 3’-end.

Small nuclear ribonucleoproteins (snRNPs) are fundamental spliceosome subunits that sequentially bind to the pre-mRNA and associated proteins within the spliceosome complex (Matera & Wang, 2014). The U2 snRNP associates early with the 5’ and 3’ splice sites of the nascent intron in the pre-spliceosome complex. The U5 and U6 snRNPs subsequently bind to the pre-spliceosome complex during assembly of the pre-catalytic spliceosome. The multisubunit Prp19 complex or NineTeen Complex (Prp19C/NTC) is incorporated as the pre-catalytic spliceosome matures to the activated spliceosome (Chan et al., 2003). The U2, U5, and U6 snRNPs and Prp19C/NTC remain part of the spliceosome as it proceeds through the splicing cycle (Frendewey & Keller, 1985; Hogg et al., 2010). At the completion of splicing, these snRNPs and the Prp19C/NTC stay bound to the intron lariat in an assembly called the intron lariat spliceosome complex (Wan et al., 2017). Disassembly of the intron lariat spliceosome complex releases the intron lariat for ensuing degradation and allows recycling of the snRNPs, Prp19C/NTC, and other spliceosome subunits into subsequent spliceosomes (Arenas & Abelson, 1997; Martin et al., 2002).

Efficient removal of intron lariats after their release from the spliceosome requires linearization, or debranching, through removal of the 2’-5’ phosphodiester-bond forming the circular portion of the intron lariat (Arenas & Hurwitz, 1987; Ruskin & Green, 1985). Ribonucleases of the highly conserved RNA DEBRANCHING ENZYME 1 (DBR1) family perform this debranching function (Chapman & Boeke, 1991). DBR1 enzymes occur as single copy genes across eukaryotes from yeast to animals to plants (Ooi et al., 2001). DBR1 is necessary to organisms with a large proportion of intron-containing pre-mRNAs. DBR1 is nonessential for the budding yeast *Saccharomyces cerevisiae*, which has few transcripts with introns, while the more intron dense fission yeast *Schizosaccharomyces pombe* requires DBR1 for normal growth rate and cell morphology (Nam et al., 1997). DBR1 is essential in higher eukaryotes including mice (Findlay et al., 2014; Zheng et al., 2015), nematodes (Piano et al., 2002), and plants (Wang et al., 2004). Further, human DBR1 activity is linked to several human diseases (Mohanta & Chakrabarti, 2021), including human immunodeficiency virus infection (Ye et al., 2005), amyotrophic lateral sclerosis (Armakola et al., 2012), and cancer (Han et al., 2017).

Recent work demonstrates high levels of intron lariats interfere with microRNA (miRNA) biogenesis in *Arabidopsis thaliana* (Li et al., 2016). In the weak *dbr1-2* allele, pre-mRNA-derived intron lariats are more stable and accumulate to higher levels than normal. These intron lariats act as competitive inhibitors of primary microRNA (pri-miRNA) processing by binding to the core components of the dicer complex DICER LIKE 1 (DCL1) and HYPONASTIC LEAVES 1 (HYL1). In this way, intron lariats inhibit DCL1/HY1-mediated processing of pri-miRNAs into miRNAs. Lower miRNA production in *dbr1-2* causes pleiotropic developmental phenotypes including curly serrated leaves, increased branching, short stature, and reduced fertility (Li et al., 2016).

Mutations in the Arabidopsis *SICKLE* (*SIC)* gene alter the processing of both pri-miRNAs and pre-mRNAs through unknown mechanisms. The strong *sic-1* mutant inhibits pri-miRNA processing, reducing the biogenesis of several miRNAs (Zhan et al., 2012). *sic-1* and the weak *sic-3* allele both elevate alternative splicing of many transcripts (Marshall et al., 2016; Zhan et al., 2012), which is exacerbated by low temperatures (Marshall et al., 2016). The RNA processing defects in *sic* mutants are accompanied by a spectrum of phenotypes, including lengthening of circadian clock period, temperature-dependent circadian clock arrhythmia, elevated sensitivity to abiotic stress, and altered growth and development apparent as curly serrated leaves, increased branching, short stature, and inhibited root growth (Karampelias et al., 2016; Marshall et al., 2016; Zhan et al., 2012).

The SIC protein is a 35 kDa proline-rich nuclear-localized protein that is conserved across the Angiosperm lineage (Marshall et al., 2016; Zhan et al., 2012). The molecular function of SIC is unknown. Most of the SIC amino acid sequence is unique except for a highly conserved MPLKIP motif (Marshall et al., 2016). This motif was first identified in human M-phase-specific PLK1-interacting protein (MPLKIP), which is associated with the human disease trichothiodystrophy non-photosensitive type 1 (NP-TTDN1) (Nakabayashi et al., 2005; Zhang et al., 2007). The core MPLKIP amino acid sequence, S M L/A E D P W, occurs in fungal, animal, and plant proteins (InterPro family IPR028265; (Paysan-Lafosse et al., 2023)). Recent work shows TTDN1 promotes DBR1 interaction with the intron-binding complex, a key component of the spliceosome, which involves TTDN1 binding of DBR1 through its MPLKIP motif (Townley et al., 2023). Further, mice deficient in *ttdn1* exhibit intron lariat accumulation and aspects of NP-TTDN1 disease.

Here we demonstrate *sic-1* and *sic-3* accumulate many of the same intron lariats as *dbr1-2* and *dbr1-3*, a newly characterized viable *dbr1* allele. Computational analysis of *sic-3* RNAseq profiling identified 1,870 intron sequences with differential usage in *sic-3*. These sequences have characteristics of intron lariats instead of annotated splice variants. Direct molecular analysis of selected differential usage intron sequences in *sic-3* and *sic-1* confirms these are intron lariats and that these RNA species also occur in *dbr1-3*. Furthermore, many of the predicted intron lariats identified in *sic-3* match confirmed intron lariats found previously in *dbr1-2.* Thus, *sic* mutants are like *dbr1* mutants because they accumulate intron lariats. Genetic analysis indicated a potential synthetic lethal interaction between *sic-1* and *dbr1-3*, while the *sic-3 dbr1-3* mutant indicated *sic-3* is largely epistatic to *dbr1-3*. Consistent with *SIC* and *DBR1* contributing to the same processes, the *sic*, *dbr1,* and *sic dbr1* mutants share physiological phenotypes, including inhibited root growth, limited rosette expansion, and a slower, less precise, circadian clock. Finally, we show the MPLKIP amino acid motif in SIC mediates physical interaction with DBR1*in vitro* and is required for SIC activity *in vivo*. These finding show a conserved role for MPLKIP containing proteins like SIC is interaction with cognate DBR1 proteins to promote debranching of intron lariats.

## RESULTS

### *sic-3* causes global changes in intron levels and transcript splice variants

Previous work showed that *sic-3* accumulates high levels of splice variants for several circadian clock-associated transcripts, a phenotype exacerbated by exposure to 16°C (Marshall et al., 2016). To understand the global impact of *sic-3* on transcript splicing at 16°C, we performed the RNAseq experiment described in SFigure 1A (Marshall & Harmon, 2022) and calculated differential usage of exons or introns in *sic-3* relative to wild-type Columbia-0 (Col-0) with the ASpli computational pipeline (Mancini et al., 2021) (Supplementary Methods). The ASpli approach divides genes into sub-genic exon and intron features based on existing annotations and then tests for differential usage of these features. The ASpli pipeline assigns features to four common splice variant types (SFigure 1B): intron retention (IR), exon skip (ES), alternative 5’ splice site (alt5ss), and alternative 3’ splice site (alt3ss). Features not matching these canonical types are reported as type “undefined”. Features with differential usage in *sic-3* were those with a log2 fold change increase >1 and a Benjamini-Hochberg (Benjamini & Hochberg, 1995) corrected false discover rate (FDR) value ≤0.05.

According to this analysis, a total of 1,968 ASpli features exhibit differential usage in *sic-3* relative to Col-0 (STable 3). Common splice variants accounted for 2.4% (47 total) of differential usage features, while 97.6% (1,921 total) were classified as the undefined type (SFigure 1C). Of the common splice variant types, IR (18 total) and ES (16 total) were most frequent, followed by alt5ss (9 total) and alt3ss (4 total) (SFigure 1C). Notably, 1,891 (98.4%) of the undefined type features corresponded to intron sequences (STable 3). To understand what transcript changes the undefined events represented, we examined the read alignment patterns for six randomly selected differential usage splice variants derived from introns taken from across the fold-change spectrum (Figure 1A). Visual comparison of the read alignment patterns at *AT1G30470 intron 1* (*I1*), *AT1G04950 I5*, *AT3G12140 I1*, *AT4G19490 I10*, *AT5G45140 I15*, and *AT2G30520 I2* revealed several attributes in common for *sic-3* and distinct from Col-0 (Figure 1B-G). These attributes included an abrupt increase in reads beginning at or close to the 5’-end of the intron (Figure 1B-G), a steep drop-off in reads at a 3’ position distinct from the 3’-end of the intron, and few or no reads overlapping the upstream or downstream exon-intron junctions. These features are unlike known transcript splice variants. However, intron lariats were discovered to have these characteristics in RNAseq profiling experiments of several human tissues (Taggart & Fairbrother, 2018). Thus, some proportion of the undefined differential usage features in *sic-3* potentially were derived from intron lariats.

**Figure 1.**
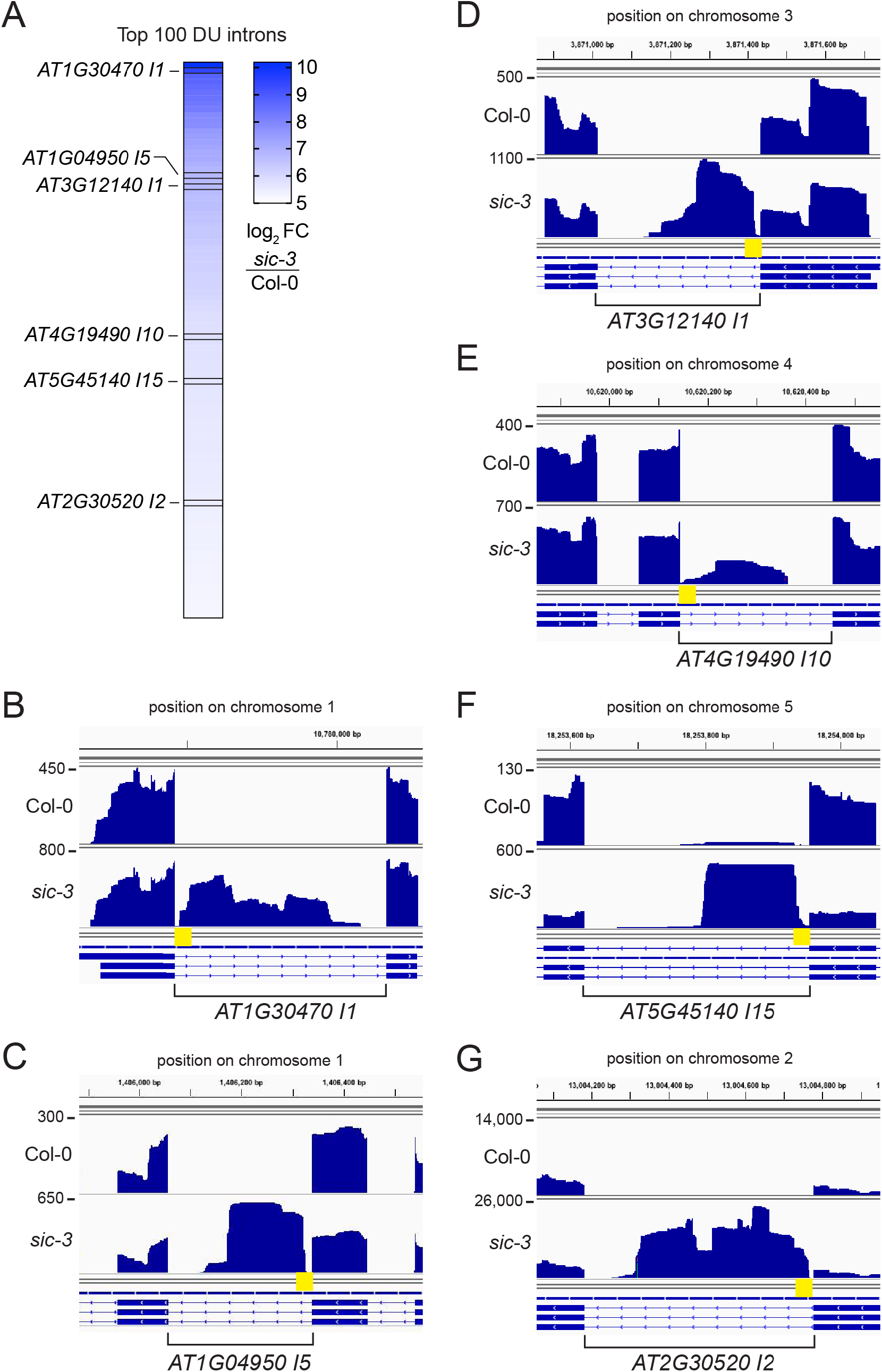
*sic-3* accumulates unusual sequences from introns. A) Top 100 introns with the greatest log_2_ fold-change (FC) increase called as undefined differential usage (DU) introns in *sic-3* by the ASpli analysis pipeline. Boxes and labels highlight introns investigated in follow-up studies. B-D) RNAseq read alignment profiles for introns *AT1G30470 I1* (B), *AT1G04950 I5* (C), *AT3G12140 I1* (D), *AT4G19490 I10* (E), *AT5G45140 I15* (F) and *AT2G30520 I2* (G) in Col-0 (top panel) and *sic-3* (bottom panel). Scale is read count and yellow box highlights intron 5’-end. Images were generated by the Integrative Genomics Viewer, version 2.4.1. (Robinson et al., 2011).

### Excess intron sequences in *sic-3* are intron lariats

To investigate potential intron lariats accumulation in *sic-3*, we employed a computational approach modeled on the ShapeShifter method previously developed for a meta-analysis of human RNAseq samples to identify intron lariat-derived sequences (Taggart & Fairbrother, 2018). ShapeShifter employs K-means clustering of RNAseq read pileup patterns to find introns with the abrupt increase in reads close to the 5’-end of an intron generated by sequencing of an intron lariat, a pattern like that displayed by the undefined differential usage introns observed in *sic-3* (Figure 1B-G).

ShapeShifter analysis was carried out as described in SFigure 2A and Supplementary Methods. The analysis pipeline was run independently for each of the three *sic-3* RNAseq replicates. Criteria applied to include introns in the ShapeShifter pipeline were intron size ≥100 base pairs and ≥10 reads aligned to positions 10-110 base pairs downstream of the intron start (Figure 2A). This analysis binned introns into five K-means clusters (STable 4), each with a distinct read accumulation pattern (Figure 2B). Independent analysis of each RNAseq replicate yielded the same read accumulation patterns. The five consensus K-means clusters representing introns shared between all three replicates contained a total of 2,576 introns (STable 5).

**Figure 2.**
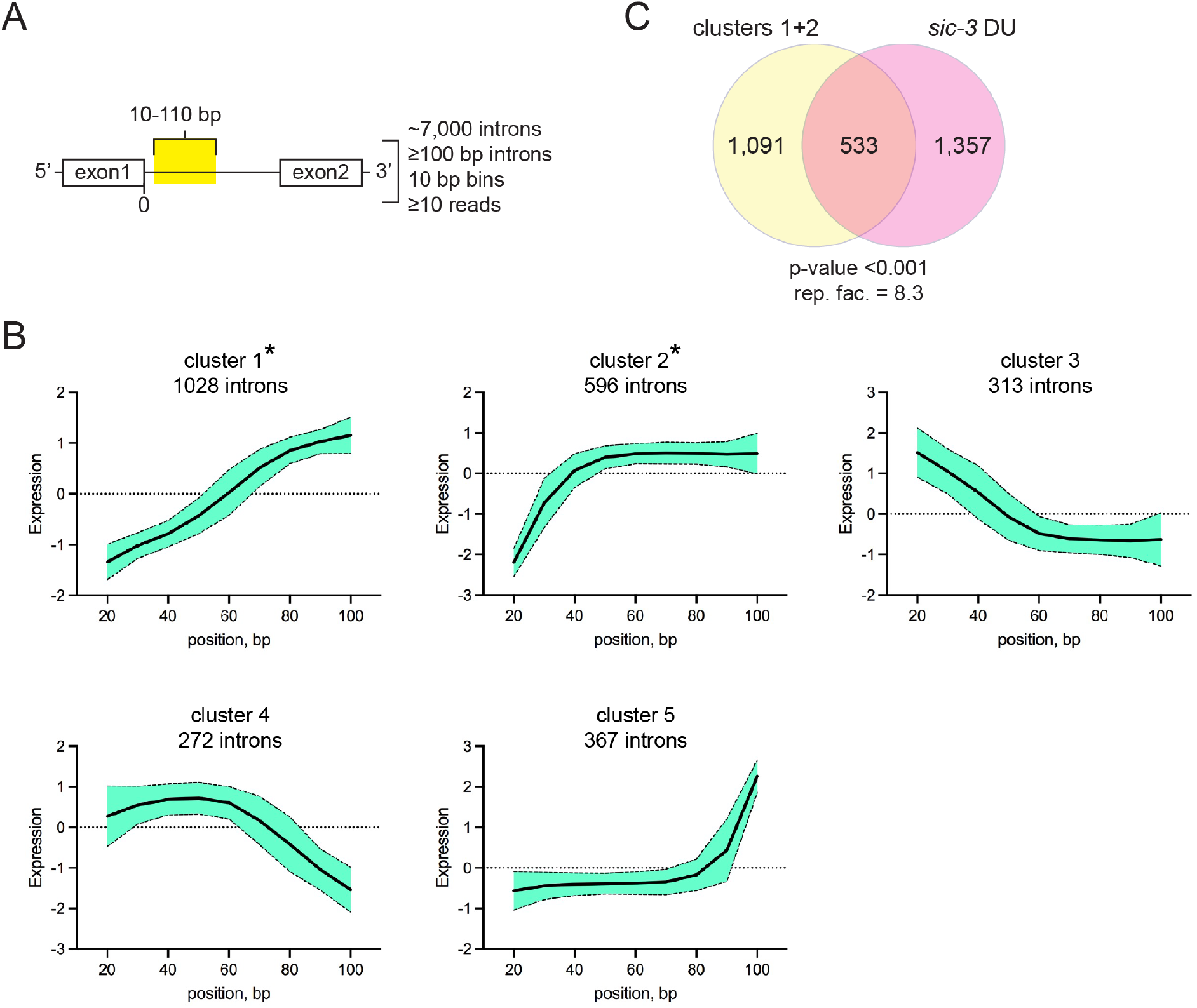
Intron sequences in *sic-3* have features of intron lariats. A) Intron features assessed by the ShapeShifter analysis pipeline. B) Consensus ShapeShifter generated clusters of *sic-3* expressed introns. Asterisk (*) marks clusters 1 and 2 containing most *sic-3* undefined differential usage introns. Shading around lines indicates standard deviation. C) Shared introns between ShapeShifter clusters 1 and 2 (clusters 1+2) and ASpli *sic-3* undefined differential usage introns (*sic-3* DU). Significance testing by hypergeometric probability at p-value <0.001 and representation factor (rep. fac.), where value >1 indicates more overlap than expected for two independent groups.

Clusters 1 and 2 represented 1,624 introns, and these corresponded to cases where read alignment rose steeply beginning within 20-60 base pairs of the intron 5’-end (Figure 2B). These two clusters contained the majority (64.7%) of *sic-3* undefined differential usage introns placed into clusters by the ShapeShifter analysis (SFigure 2B). Clusters 1 and 2 together shared 533 introns (352 in cluster 1 and 181 in cluster 2) with the 1,890 undefined differential usage introns called by ASpli (Figure 2C), a representation factor of 8.3 for all introns, and a statistically significant enrichment (hypergeometric probability p-value <0.001) (Supplementary Methods). Included in clusters 1 and 2 were the six randomly selected differential usage intron sequences examined above: *AT1G30470 I1, AT1G04950 I5*, *AT3G12140 I1*, *AT4G19490 I10*, *AT5G45140 I15*, and *AT2G30520 I2*. The ShapeShifter analysis supported the notion that many of the undefined differential usage intron sequences in *sic-3* come from intron lariats.

We employed reverse transcription PCR (RT-PCR) to confirm the presence of intron lariats in *sic-3* and quantitative RT-PCR (qPCR) to quantify intron lariat levels. In standard cDNA synthesis reactions primed with random primer, intron lariats appear as a unique juxtaposition of sequences from the intron 5’-end and the lariat branchpoint, because MMLV reverse transcriptase reads through the 2’-5’ bond at the branchpoint (Vogel et al., 1997) (Figure 3A). PCR with primers flanking this unique junction selectively amplify the target intron lariat. To increase the stringency of RT-PCR tests, total RNA was treated with *Escherichia coli* RNase R before cDNA synthesis (Figure 3A). RNase R is a single-strand RNA-specific nuclease that degrades linear RNAs, like mRNA and ribosomal RNA, but does not cleave circular RNAs, such as the circular portion of intron lariats (Suzuki et al., 2006; Vincent & Deutscher, 2006). Prior treatment of *sic-3* and Col-0 total RNA samples with RNase R reduced by at least 120-fold the cDNA levels from the *IPP2* transcript (SFigure 3), a control transcript for RT-PCR and qPCR with stable and constitutive expression (Hazen et al., 2005).

**Figure 3.**
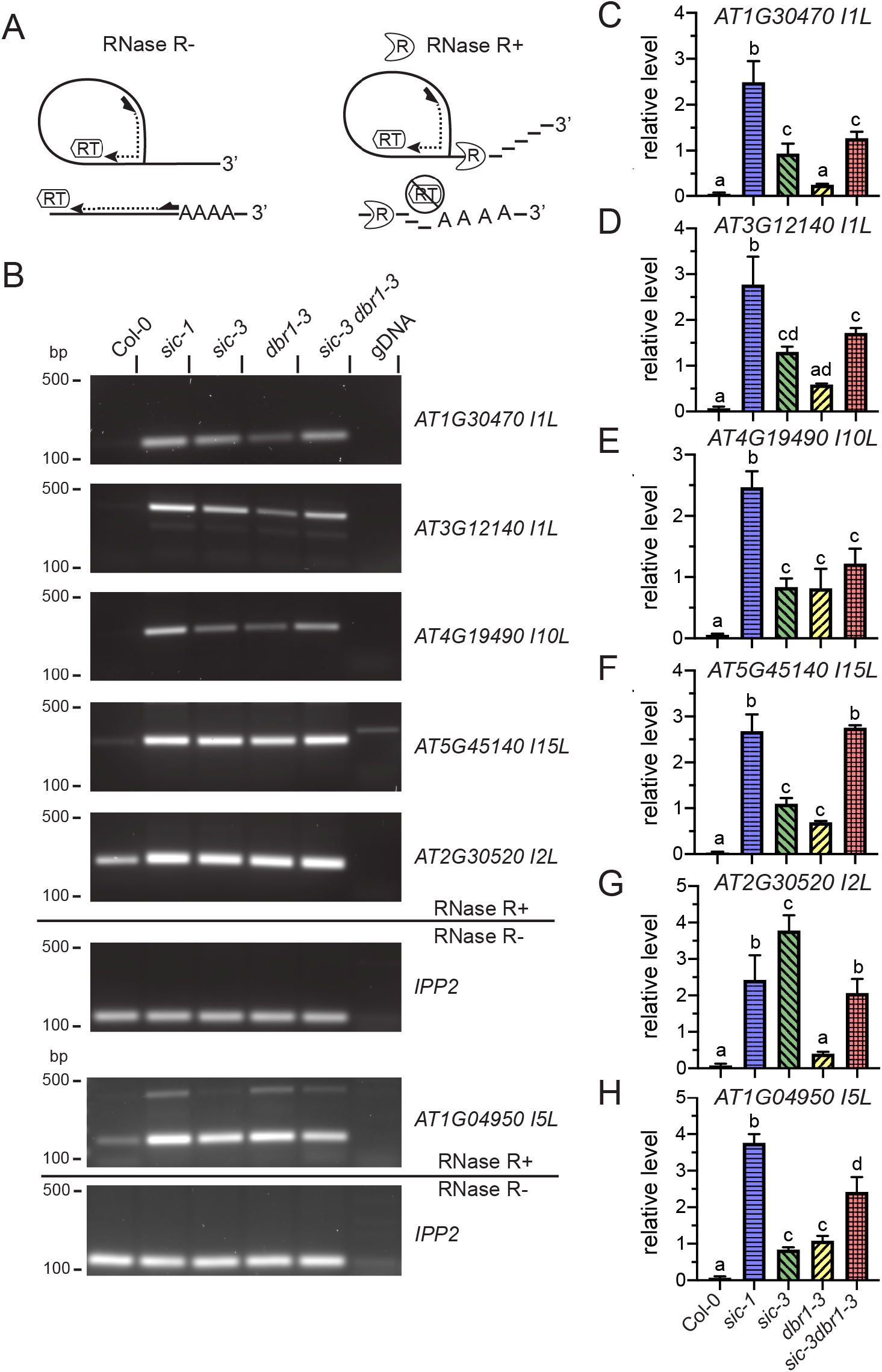
Intron sequences in *sic-3*, *dbr1-3,* and *sic-3 dbr1-3* are from intron lariats. A) RNase R degradation of mRNAs enriches for synthesis of intron lariat cDNAs. Shapes indicate enzymes reverse transcriptase (RT) and RNase R (R). Short thick arrows are random primers and dotted lines with arrows are cDNAs. B) Ethidium bromide-stained RT-PCR products for intron lariats *AT1G30470 I1L*, *AT3G12140 I1L*, *AT4G19490 I10L*, *AT5G45140 I15L*, *AT2G30520 I2L*, and *AT1G04950 I5L* from RNase R treated samples (RNase R+) prepared from the indicated genotypes. Genomic DNA from Col-0 (gDNA) was a negative control template. The loading control was *IPP2* transcript amplified from samples not treated with RNase R (RNase R-). All samples were run out on the same agarose gel, except *AT1G09450 I4L* and the corresponding *IPP2* control. Position of 500 and 100 base pair (bp) standards indicated at left. Images are representative of three independent biological replicates. C-H) Quantification of intron lariats C) *AT1G30470 I1L*, D) *AT3G12140 I1L* E), *AT4G19490 I10L*, F) *AT5G45140 I15L*, G) *AT2G30520 I2L*, and H) *AT1G04950 I5L* by qPCR from RNase R treated samples of Col-0 (white solid bar), *sic-1* (blue horizontal hatched bar), *sic-3* (green left crosshatched bar), *dbr1-3* (yellow right crosshatched bar), and *sic-3 dbr1-3* (red checkered bar). The error bars are standard deviation of three independent biological replicates. Means sharing a common letter are not significantly different by one-way ANOVA with Tukey’s multiple comparisons test at p <0.05 level of significance.

The six introns examined above (Figure 1B-G) were tested for the intron lariat sequence configuration in seedlings of Col-0, *sic-3*, and the strong *sic-1* mutant allele (Zhan et al., 2012). RT-PCR amplified DNA products of the expected size from *sic-3* and *sic-1* cDNA made from RNase R-treated total RNA for *AT1G30470 intron 1 lariat* (*I1L*), *AT3G12140 I1L*, *AT4G19490 I10L*, *AT5G45140 I15L*, *AT2G30520 I2L*, and *AT1G04950 I5L*, but not when Col-0 genomic DNA (gDNA) was provided as a template (Figure 3B). A clear band was not amplified from Col-0 cDNA for *AT1G30470 I1L*, *AT3G12140 I1L*, *AT4G19490 I10L*, and *AT5G45140 I15L*, while a lower intensity band matching the intron lariat band was amplified for *AT2G30520 I2L* and *AT1G04950 I4L* with this template.

Consistent with the templates of these RT-PCR products arising from intron lariats, the sequence of each RT-PCR product amplified from *sic-3* had a structure expected for a lariat. An internal 5’-3’ sequence from near the end of the intron was joined to the 5’-end of the same intron (SFigure 4). The position of the branchpoint in *AT3G12140 I1L*, *AT2G30520 I2L*, *AT4G19490 I10L*, and *AT5G45140 I15L* matched that identified by prior RNAseq analysis (Zhang et al., 2019), while *AT1G30470 I1L* and *AT1G04950 I4L* are undescribed lariat structures.

Quantification of these test introns with the suspected intron lariat sequence configuration with qPCR revealed these species occurred in *sic-3* and *sic-1* at levels 25-fold and 50-fold higher, respectively, than in Col-0 (Figure 3C-H). Intron lariat accumulation was greater in *sic-1* than *sic-3* for *AT1G30470 I1L*, *AT3G12140 I1L*, *AT4G19490 I10L*, *AT5G45140 I15L*, and *AT1G04950 I4L* (Figure 3C-F, H), while *AT2G30520 I2L* was at higher levels in *sic-3* (Figure 3G). Since *DBR1* transcript levels in *sic-1* and *sic-3* were not different from Col-0 (SFigure 5B), the *sic* alleles themselves cause intron lariat accumulation instead of through an indirect effect on *DBR1* expression. In total these findings show *sic* mutants amass intron lariats at levels many times greater than Col-0 and many of the undefined differential usage intron sequences in *sic-3* are likely derived from intron lariats.

### Intron lariats in *sic-3* and *sic-1* match those in the weak *dbr1-3* mutant

The tendency of *sic* mutants to have abundant intron lariats is like that described previously for the weak *dbr1-2* mutant (Li et al., 2016). To enable comparisons between *dbr1* and *sic* mutant alleles, we sought a viable Arabidopsis *dbr1* mutant allele from the SALK T-DNA insertion collection (Alonso et al., 2003). One line had a T-DNA insertion in the 5’-untranslated region (UTR) of *DBR1* at nucleotide position 427, just upstream of the translation start codon (SFigure 5A). This T-DNA reduced *DBR1* expression to nearly half of normal levels (SFigure 5B). The mutant, designated *dbr1-3*, is viable as a homozygote, indicating the T-DNA does not eliminate *DBR1* function. *SIC* transcript levels in *dbr1-3* were statistically indistinguishable from Col-0 (SFigure 5C), indicating *SIC* expression is not affected by *DBR1*. The *AT1G30470 I1L*, *AT3G12140 I1L*, *AT4G19490 I10L*, *AT5G45140 I15L*, *AT2G30520 I2L*, and *AT1G04950 I5L* products were amplified by RT-PCR from *dbr1-3* cDNA made from RNase R-treated total RNA (Figure 3B). Thus, *dbr1-3* interferes with *DBR1* function sufficiently to cause intron lariat accumulation without the embryo lethality seen in the *dbr1-1* allele (Wang et al., 2004).

The intron lariat-derived RT-PCR products in *dbr1-3* matched those from *sic-3* and *sic-1* (Figure 3B). Quantification of these intron lariat sequences in *dbr1-3* indicated levels of *AT3G12140 I1L*, *AT4G19490 I10L*, *AT5G45140 I15L,* and *AT1G04950 I5L* were comparable to those in *sic-3* (Figure 3D-F, H), while *AT1G30470 I1L* and *AT2G30520 I2L* were lower than in *sic-3* and at levels statistically indistinguishable from Col-0 (Figure 3C, G). In all cases, levels of intron lariat sequences in *dbr1-3* were lower than in *sic-1*. Thus, *dbr1-3* reduces *DBR1* function, and this allele causes the accumulation of intron lariats that match those in *sic-3* and *sic-1*.

### Many of the same intron lariat sequences occur in *sic-3* and *dbr1-2*

A previous study identified differentially expressed intron lariats in *dbr1-2* based on RNAseq profiling of RNase R-treated total RNA from *dbr1-2* and Col-0 (Li et al., 2016). To determine if *sic-3* and *dbr1-2* similarly affect intron lariat accumulation, we compared the 1,300 RNase R-resistant introns from *dbr1-2* (STable 6) to the 1,891 *sic-3* unidentified differential usage introns discovered by ASpli analysis (STable 3). A total of 235 *dbr1-2* intron lariats were shared between the two sets (Figure 4A), a statistically significant enrichment (hypergeometric probability at p-value < 0.001) and a representation factor of 4.6. Both intron lists included the RT-PCR-validated intron lariats *AT3G12140 I1L*, *AT4G19490 I10L*, *AT5G45140 I15L*, and *AT2G30520 I2L* (Figure 4B). In the RNase R treated *dbr1-2* and Col-0 samples, the read alignment patterns for these four features had the characteristics of intron lariats observed in the *sic-3* RNAseq profiling (SFigure 6A-D): an abrupt increase in reads at the beginning of 5’-end of the intron, a steep drop-off in reads in the middle of the intron, and a lack of reads overlapping intron-exon junctions. The read counts for these sequences in *dbr1-2* were at least 3-fold greater than in Col-0, indicating these intron lariats normally occur in Col-0 and *dbr1-2* amplifies their accumulation (SFigure 6A-D. These findings indicate *sic-3* and *dbr1-2* cause buildup of many of the same intron lariats.

**Figure 4.**
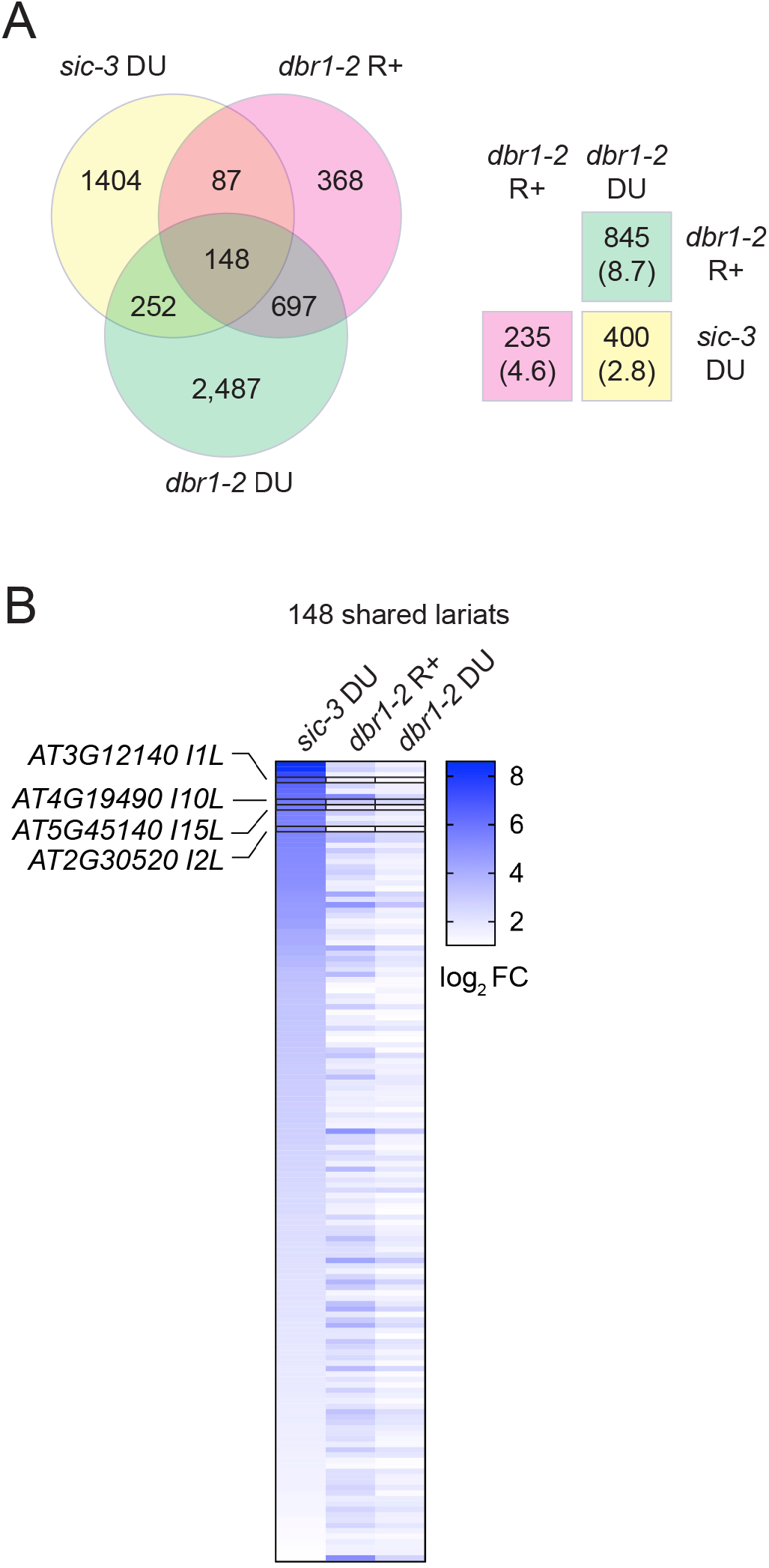
Intron sequences in *sic-3* match *dbr1-2* RNase R-resistant intron lariats and *dbr1-2* differential usage introns. A) Overlap between the 1,891 *sic-3* undefined differential usage introns from ASpli analysis (*sic-3* DU), the 1,300 *dbr1-2* RNase R-resistant intron sequences (*dbr1-2* R+) (Li et al., 2016), and the 3,584 *dbr1-2* unidentified differential usage introns from ASpli analysis (*dbr1-2* DU). Inset boxes at right indicate number of intron sequences shared between the two sets indicated. Number in parenthesis is representation factor, where a value >1 indicates more overlap than expected for two independent groups. Overlap between all sets is significant by hypergeometric probability testing at p-value <0.001. B) Log_2_ fold-changes relative to Col-0 for the 148 introns in common for the *sic-3* DU, *dbr1-2* R+, and *dbr1-2* differential usage sets. Labels and boxes highlight introns investigated in follow-up studies.

To determine if the *dbr1-2* transcriptome exhibits an overabundance of intron sequences like the *sic-3* transcriptome, we analyzed an existing RNAseq profiling dataset of *dbr1-2* and Col-0 without RNase R treatment (Li et al., 2016) with the ASpli pipeline to identify splice variants with differential usage in *dbr1-2* (STable 7). This analysis indicated 3,584 (93.3%) of differential usage splice variants in *dbr1-2* were intron sequences of the undefined type (SFigure 6E). This bias in ASpli called features is like that observed in *sic-3* (SFigure 1C). A total of 400 of the *dbr1-2* sequences were shared with the *sic-3* unidentified differential usage intron set (Figure 4A), which is a statistically significant enrichment (hypergeometric probability at p-value < 0.001) and a representation factor of 2.8. The shared set includes RT-PCR-validated intron lariats *AT3G12140 I1L*, *AT4G19490 I10L*, *AT5G45140 I15L*, and *AT2G30520 I2L* (Figure 4B). In addition, a total of 845 introns in the *dbr1-2* differential usage set were shared with the 1,300 RNase R resistant introns in *dbr1-2* (Figure 4A), a statistically significant enrichment (hypergeometric probability at p-value < 0.001) and a representation factor of 8.7. Thus, intron sequences derived from intron lariats occur at high levels in *sic-3* and *dbr1-2*. We conclude *sic* mutants interfere with a key aspect of intron lariat processing and this leads to intron lariat accumulation comparable to when *DBR1* activity is diminished as in *dbr1-2* and *dbr1-3*.

### Testing genetic interaction between *sic* alleles and *dbr1-3*

We sought to combine *dbr1-3* with *sic-1* and *sic-3* to gain information on the genetic relationship between *SIC* and *DBR1*. To generate a *sic-1 dbr1-3* mutant, homozygous *sic-1* (*sic-1*/*sic-1*) emasculated flowers were fertilized with homozygous *dbr1-3* (*dbr1-3*/*dbr1-3*) pollen. Plants heterozygous for *sic-1* (+/*sic-1*) and homozygous for *dbr1-3* (*dbr1-3*/*dbr1-3*) were identified in the F2 progeny and these were allowed to self. A total of 112 progeny from this selfing were scored for *sic-1* and *dbr1-3* with the expectation of recovering at least 11 *sic-1*/*sic-1 dbr1-3*/*dbr1-3* plants (probability = 95%). None of the plants in this family were *sic-1*/*sic-1 dbr1-3*/*dbr1-3*, while 83 plants (74%) were +/*sic-1 dbr1-3*/*dbr1-3* and 29 (26%) were +/+ *dbr1-3*/*dbr1-3* (Table 1). Chi-square and Fisher’s exact testing indicated the observed segregation significantly deviated from the expected Mendelian ratio of 1:2:1 for the recessive *sic-1* allele (p-value <0.005, Table 1). Thus, the *sic-1 dbr1-3* mutant was difficult to recover, indicating this combination of alleles may cause a synthetic lethal phenotype.

**Table 1.** Outcome of *sic-1* X *dbr1-3* and *sic-3* X *dbr1-3* crosses.

| Cross<sup>a</sup> | Expected Genotypes | Number Recovered/ Expected<sup>b</sup> | Percent Recovery/ Expected<sup>c</sup> |
| --- | --- | --- | --- |
| <u>*+/sic-1 dbr1-3/dbr1-3* ♀</u> |  |  |  |
| <u>*+/sic-1 dbr1-3/dbr1-3* ♂</u> |  |  |  |
|  | *sic-1/sic-1 dbr1-3/dbr1-3* | 0/11 | 0/25 |
|  | *+/sic-1 dbr1-3/dbr1-3* | 83/26 | 74/50 |
|  | *+/+ dbr1-3/dbr1-3* | 29/11 | 26/25 |
| Chi-square value<sup>d</sup> (p-value) |  | 33.262<br>(5.987e-08) |  |
| <u>*sic-3/sic-3 +/dbr1-3* ♀</u> |  |  |  |
| <u>*+/sic-3 dbr1-3/dbr1-3* ♂</u> |  |  |  |
|  | *sic-3/sic-3 dbr1-3/dbr1-3* | 9/6 | 14/25 |
|  | *sic-3/sic-3 +/dbr1-3* | 21/6 | 32/25 |
|  | *+/sic-3 +/dbr1-3* | 17/6 | 26/25 |
|  | *+/sic-3 dbr1-3/dbr1-3* | 18/6 | 28/25 |
| Chi-square value<sup>d</sup> (p-value) |  | 2.794<br>(0.4245) |  |

<sup>a</sup> ♀ - female; ♂ - male
<sup>b</sup>At probability = 95%.
<sup>c</sup>Of total plants tested.
<sup>d</sup>See Supplementary Methods for details.

To obtain the homozygous *sic-3 dbr1-3* mutant, *sic-3*/*sic-3* flowers were pollinated with *dbr1-3*/*dbr1-3* pollen and then F2 generation *sic-3*/*sic-3 +/dbr1-3* emasculated flowers were pollinated with +/*sic-3 dbr1-3*/*dbr1-3* pollen. Testing of 65 progeny from this cross identified all the expected genotypes (Table 1). Individuals of the +/*sic-3* +/*dbr1-3* and +/*sic-3 dbr1-3*/*dbr1-3* genotypes appeared at close to the expected frequency of 25%, while the *sic-3*/*sic-3 dbr1-3*/*dbr1-3* genotype was less frequent (14%) and the *sic-3*/*sic-3* +/*dbr1-3* genotype was more frequent (32%). Chi-square and Fisher’s exact testing indicated no segregation distortion occurred in this case (p-value <0.05, Table 1).

The intron lariat phenotype of *sic-3 dbr1-3* seedlings was like *sic-3* and *dbr1-3*. With cDNA made from *sic-3 dbr1-3* RNase R-treated RNA as a RT-PCR template, the primers targeting the intron lariats investigated above amplified *AT1G30470 I1L*, *AT3G12140 I1L*, *AT4G19490 I10L*, *AT5G45140 I15L*, *AT2G30520 I2L*, and *AT1G04950 I5L* (Figure 3B). qPCR quantification of these species in *sic-3 dbr1-3* seedlings revealed higher accumulation relative to *dbr1-3* (Figure 3C-H). The exception was *AT4G19490 I10L*, where levels were indistinguishable between *dbr1-3*, *sic-3*, and *sic-3 dbr1-3* (Figure 3E). Levels of these intron lariats were comparable to or higher than in *sic-3*, but lower than in *sic-1*, for all except *AT2G30520 I2L*, where *sic-3* had the highest levels of the tested genotypes (Figure 3G), and *AT5G45140 I15L*, where levels were indistinguishable between *sic-3 dbr1-3* and *sic-1* (Figure 3F). The results indicate *sic-3* is largely epistatic to *dbr1-3*.

### Disrupted growth and circadian clock activity in *sic-3*, *dbr1-3*, and *sic-3 dbr1-3*

To determine if intron lariat accumulation causes a common set of changes in plant physiological processes, we assessed root growth in seedlings and rosette expansion in adult plants of *sic-3*, *dbr1-3*, and *sic-3 dbr1-3*. These aspects of plant growth have been reported to be altered in *sic* mutant (Karampelias et al., 2016; Marshall et al., 2016; Zhan et al., 2012) and *dbr1-2* (Li et al., 2016). Consistent with these previous findings, the mean root lengths of 7 day-old *sic-3* and *dbr1-3* seedlings were shorter than the mean of the Col-0 seedlings (Figure 5A, B). *sic-3* showed greater root growth inhibition than *dbr1-3* (Figure 5B). The roots of *sic-3 dbr1-3* seedlings were also shorter than Col-0 and the mean length was between that of *sic-3* and *dbr1-3*. Adult rosettes of *sic-3* and *dbr1-3* were smaller in diameter than Col-0 and the magnitude of this effect was similar for each allele (Figure 5 C, D). While smaller than Col-0, the size of *sic-3 dbr1-3* rosettes were indistinguishable from *sic-3* (Figure 5D). Thus, *sic-3* and *dbr1-3* individually inhibited root growth and rosette expansion and there was no additive effect when these two alleles were combined.

**Figure 5.**
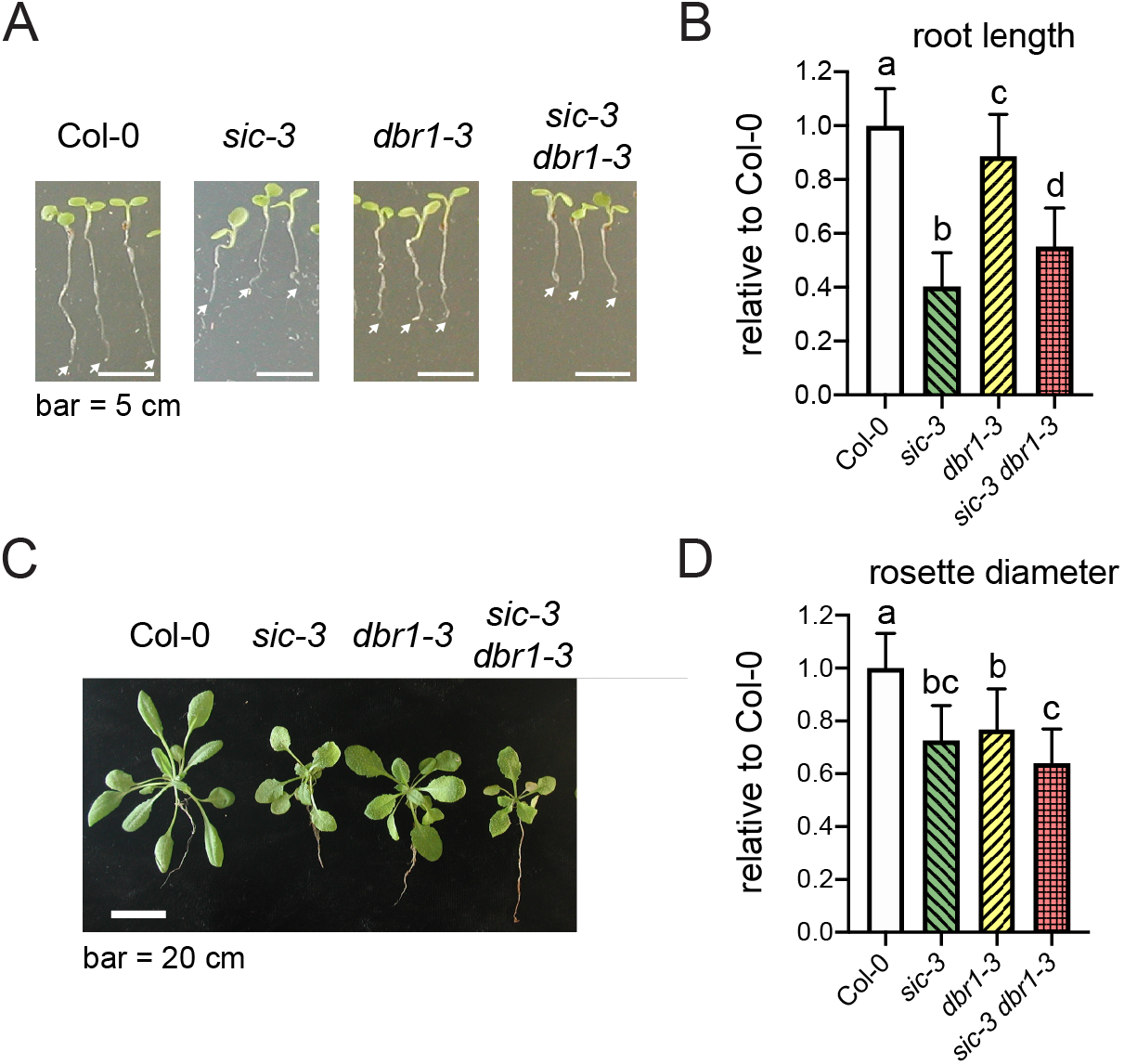
Growth inhibition caused by *sic-3*, *dbr1-3*, and *sic-3 dbr1-3*. A) Representative 7-day-old seedlings of the indicated genotype. White arrows indicate root tip. Bar is 5 cm. B) Mean root length of Col-0 (white solid bar), *sic-3* (green left crosshatched bar), *dbr1-3* (yellow right crosshatched bar), and *sic-3 dbr1-3* (red checkered bar) seedlings (n = 28 for all genotypes). The value for each seedling was normalized to the mean for Col-0. Data from 2 independent biological replicates. Error bars are standard deviation. C) Representative rosettes of 20-day-old plants of the indicated genotype. Bar is 20 cm. D) Mean rosette diameter for Col-0 (white solid bar; n = 10), *sic-3* (green left crosshatched bar; n = 16), *dbr1-3* (yellow right crosshatched bar; n = 22), and *sic-3 dbr1-3* (red checkered bar, n = 22) plants from two independent biological replicates. The value for individual were normalized to the mean of Col-0. Error bars are standard deviations. Means sharing a common letter are not significantly different by one-way ANOVA with Tukey’s multiple comparisons test at p <0.05 level of significance.

We assessed the effect of *sic-3 dbr1-3* and *dbr1-2* on the accuracy (i.e., mean estimated period) and precision (i.e., estimated period variation in the population) of circadian clock with the *PRR7:LUC+* reporter (Salome & McClung, 2005), because *sic-3* and *sic-1* exhibit longer and less consistent periods (Marshall et al. 2016). The *sic-3 dbr1-3* population exhibited a high degree of variation in bioluminescence rhythms, which was distinct from *sic-3* and *sic-1* (Figure 6A). As a result, the mean rhythm for the *sic-3 dbr1-3* population was damped relative to all the other genotypes. The mean estimated period (±standard deviation) of *sic-3 dbr1-3* was 27.1 (±3.0) hours, which was not statistically different from the mean of 27.4 (±1.3) hours in *sic-1*, but a substantial lengthening of period from the mean of 24.4 (±0.9) hours in Col-0 (Figure 6B). Notably, the estimated periods from the *sic-3 dbr1-3* population spanned a nearly 15-hour range, indicating a substantial loss of circadian clock precision in this mutant that is even greater than that in *sic-1*. On the other hand, the mean estimated period for *dbr1-3* of 25.4 (±0.8) hours matched the 25.7 (±0.7) hour period of *sic-3* indicating *dbr1-3* slows the pace of the clock to a similar degree as *sic-3*. These findings show the *sic-3 dbr1-3* mutant has more profound alterations in circadian clock accuracy and precision than the individual *sic* and *dbr1* mutants, and a mean period like *sic-1*.

**Figure 6.**
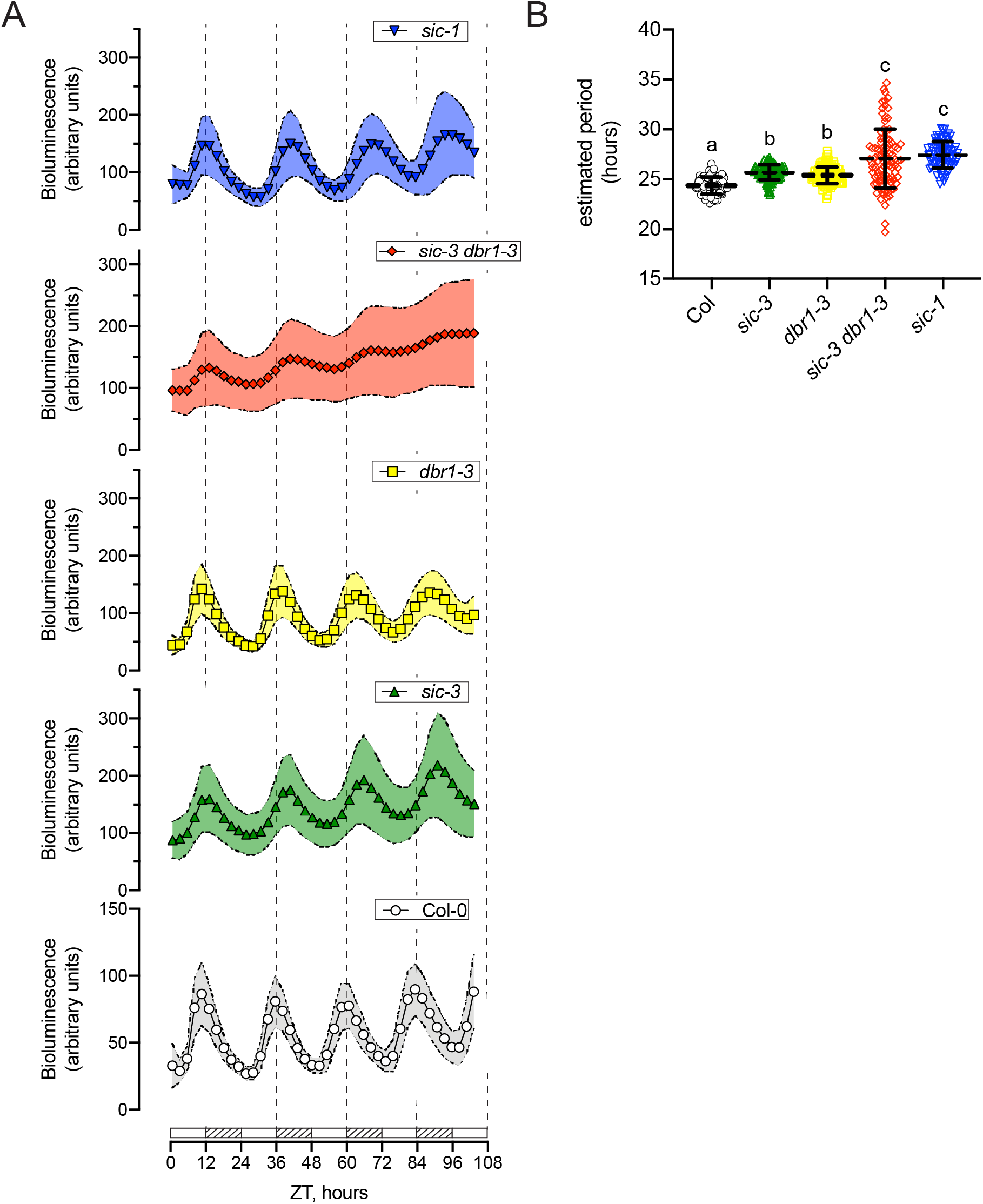
Impaired circadian clock accuracy and precision in *sic-3 dbr1-3*. Circadian rhythms (A) and estimated free-running period (B) for Col-0 (white circles; n = 91), *sic-3* (green triangles; n = 155), *dbr1-3* (yellow squares; n = 133), *sic-3 dbr1-3* (red diamonds; n = 94), and *sic-1* (blue inverted triangles; n = 76) under constant white light and 22°C. Bioluminescence is from *PRR7:LUC+* reporter (Salome & McClung, 2005). Data are from 4 independent biological replicates. A) Bioluminescence of seedlings at the indicated number of hours after transfer to constant conditions (zeitgeber time, ZT). Line and symbols indicate the mean of all seedlings of that genotype and shaded regions indicate standard deviation. White bars and hatched bars on the x-axis indicate subjective day and night, respectively. Vertical dotted lines mark the onset of subjective night. B) Estimated period of each rhythmic seedling of the indicated genotype, mean period indicated by dashed line, and error bars are standard deviation. Period estimates are from individuals with relative amplitude error values <0.6. Means sharing a common letter are not significantly different by one-way ANOVA with Tukey’s multiple comparisons test at p <0.05 level of significance.

### SIC function depends on physical interacts with DBR1 through its MPLKIP motif

A previous co-immunoprecipitation purification-mass spectrometry (CoIP-MS) study found multiple DBR1-derived peptides in a pull down of SIC protein from Arabidopsis cell suspension cultures (Karampelias et al., 2016), indicating SIC potentially interacts with DBR1 either through direct protein-protein contacts or indirectly as participants in a larger protein complex. We tested for a direct interaction between SIC and DBR1with yeast two hybrid and co-immunoprecipitation (CoIP) of *in vitro* expressed proteins.

In yeast two-hybrid, a combination of a GAL4-DNA binding domain-DBR1 fusion (DB-DBR1) bait and a GAL4 Activation Domain-SIC fusion (AD-SIC) prey allowed growth of adenine auxotrophic yeast on selective media lacking this amino acid (Figure 7A), indicating activation of the GAL4 promoter-ADE reporter construct. Under the same selective conditions, growth did not occur for either a combination of the DB-DBR1 bait with a negative control AD-human Lamin (AD-LAM) prey or the DB-LAM bait together with the AD-SIC prey (Figure 7A). Thus, SIC and DBR1 interact directly in yeast.

**Figure 7.**
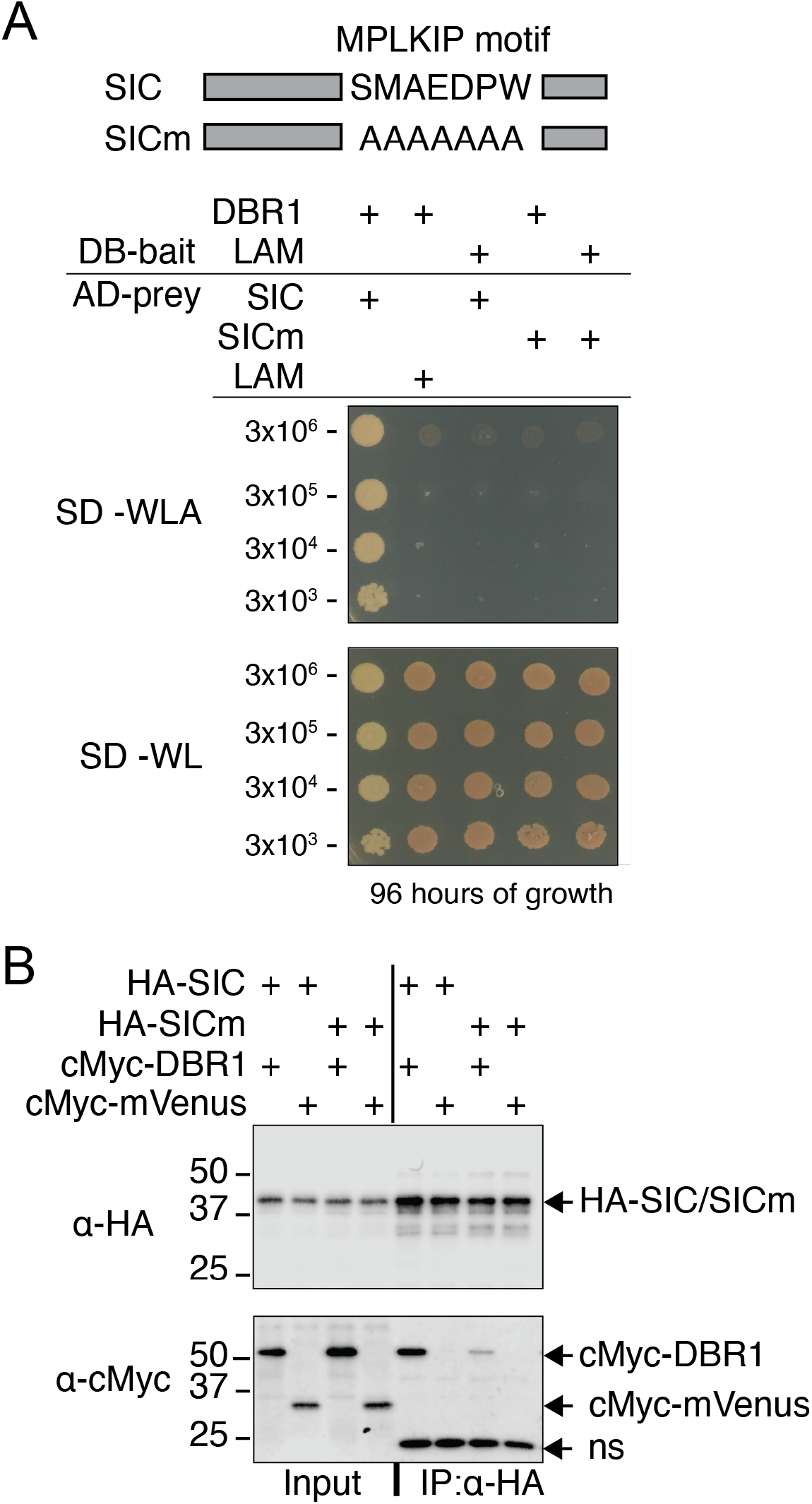
SIC physically interacts with DBR1 via its MPLKIP motif. A) Yeast two-hybrid test for interaction between the DB-DBR1 bait and AD-SIC or AD-SICm preys. Negative controls are DB-LAM and AD-LAM. Selective media was SD without W, L, and A (SD-WLA) and control media was SD without W and L (SD-WL). Cell number in spots is shown at the left of plate pictures, growth is after 96 hours at 30°C. Cartoon at top indicates the amino acid changes to the MPLKIP motif in SICm. Data are representative of three independent bait-prey cultures and three independent experiments. B) Western blot of proteins from CoIP experiments with HA-SIC or HA-SICm baits and cMyc-DBR1 or cMyc-mVenus preys. Left portion of panel are the indicated inputs to the CoIP (Input), which was the HA-tagged protein bound to magnetic anti-HA beads mixed with the indicated prey protein, Right portion is proteins isolated by the pull down (IP:α-HA). Top panel is blot probed with anti-HA as primary antibody (α-HA) and bottom panel is blot probed with anti-cMyc antibody (α-cMyc). Both were probed with anti-mouse-HRP antibody as secondary antibody for detection of bands by chemiluminescence and a Li-Cor Odyssey XF molecular imager. Arrows at the right indicate position of each protein and “ns” indicates a nonspecific protein from the pull down with anti-HA beads. Data are representative of two independent CoIP experiments.

CoIP of *in vitro* expressed proteins confirmed that SIC directly interacts with DBR1. *In vitro* transcription-translation reactions generated HA-epitope tagged SIC (HA-SIC) bait and prey proteins cMyc-epitope tagged DBR1 (cMyc-DBR1) and mVenus (cMyc-mVenus). In three independent experiments, the cMyc-DBR1 prey co-purified with the HA-SIC bait in a pull down with anti-HA magnetic beads, but the cMyc-mVenus protein did not (Figure 7B). Together, the CoIP and yeast two-hybrid experiments confirm SIC and DBR1 bind to one another, explaining co-purification of DBR1 with SIC *in vivo* (Karampelias et al., 2016).

SIC contains an MPLKIP motif with high identity to motifs present in diverse fungal, animal, and plant proteins (SFigure 7). We hypothesized the SIC -DBR1 interaction may involve the core amino acids of the SIC MPLKIP motif. To test this hypothesis, we generated the SICm construct in which the core MPLKIP amino acids S M A E D P W were substituted with A residues (Figure 7A). Unlike the wild-type AD-SIC prey in yeast two-hybrid, an AD-SICm prey expressed with the DB-DBR1 bait did not support growth on selective media lacking adenine (Figure 7A). Thus, the SICm protein lacks the capacity to interact with DBR in yeast. In two independent CoIP experiments, the HA-SICm protein exhibited diminished recovery of the cMyc-DBR1 protein relative to wild-type HA-SIC (Figure 7B). These results indicate the MPLKIP motif is important for SIC to bind to DBR1.

To confirm the MPLKIP motif contributes to SIC function *in vivo*, SIC and SICm proteins were tested for complementation of the *sic-3* lariat accumulation and circadian clock phenotypes. We constructed stable transgenic lines in *sic-3* carrying a construct made up of the SIC promoter region upstream of the genomic *SIC* sequence with a C-terminal translational fusion of fluorescent mVenus protein and the HA epitope (*SICp:SIC-V-H*) or the *SICm* genomic sequence upstream of this cassette (*SICp:SICm-V-H*). The SIC-V-H and SICm-V-H proteins were readily detectable by western blot in whole cell extracts made from these lines (SFigure 8A), Fluorescent confocal microscopy of cotyledons demonstrated both the SIC-V-H and SICm-V-H proteins accumulated in structures consistent with nuclei (SFigure 8B), as reported before for a SIC-YFP fusion (Marshall et al., 2016). Therefore, elimination of the MPLKIP motif in the SICm-V-H protein did not interfere with its targeting to the nucleus.

Expression of SIC-V-H, but not SICm-V-H, returned levels of the *AT1G30470 I1L*, *AT3G12140 I1L*, *AT4G19490 I10L*, *AT5G45140 I15L*, *AT2G30520 I2L*, and *AT1G04950* intron lariats to that observed in Col-0 wild-type plants (Figure 8). Intron lariats levels measured by qPCR in the two *SICp:SIC-V-H*/*sic-3* lines were indistinguishable from Col-0 and significantly reduced compared to *sic-3* (Figure 8A-F), On the other hand, the *SICp:SICm-V-H*/*sic-3* lines had intron lariats levels comparable to or greater than in *sic-3* (Figure 8A-F), even though SICm-V-H protein expression was greater than SIC-V-H in these lines (SFigure 8A). Thus, SICm-V-H protein without the core MPLKIP motif cannot fulfill the role played by SIC in intron lariat processing, likely due to a reduced DBR1 binding capacity.

The *SICp:SICm-V-H*/*sic-3* construct also did not complement the circadian clock phenotype of *sic-3* like the *SICp:SIC-V-H*/*sic-3* construct. Circadian rhythms according to the *PRR7:LUC+* reporter (Salome & McClung, 2005) in the two *SICp:SICm-V-H*/*sic-3* lines were similar to *sic-3* and appeared longer period than Col-0 (Figure 8G). On the other hand, the rhythm waveforms in the two *SICp:SIC-V-H*/*sic-3* lines matched Col-0 (Figure 8G). The mean estimated period for 32 individual seedlings was longer (±standard deviation) than Col-0 by 1.5 (±0.6) hours for each *SICp:SIC-V-H*/*sic-3* line, like the 1.2 (±0.7) hour period lengthening in *sic-3* (Figure 8H). In contrast, the mean estimated period of the two *SICp:SIC-V-H*/*sic-3* lines were within 0.2 (±0.5) hours of Col-0, indicating effective complementation of *sic-3* (Figure 8B). These results confirm the MPLKIP motif in SIC is required for activity *in vivo*.

**Figure 8.**
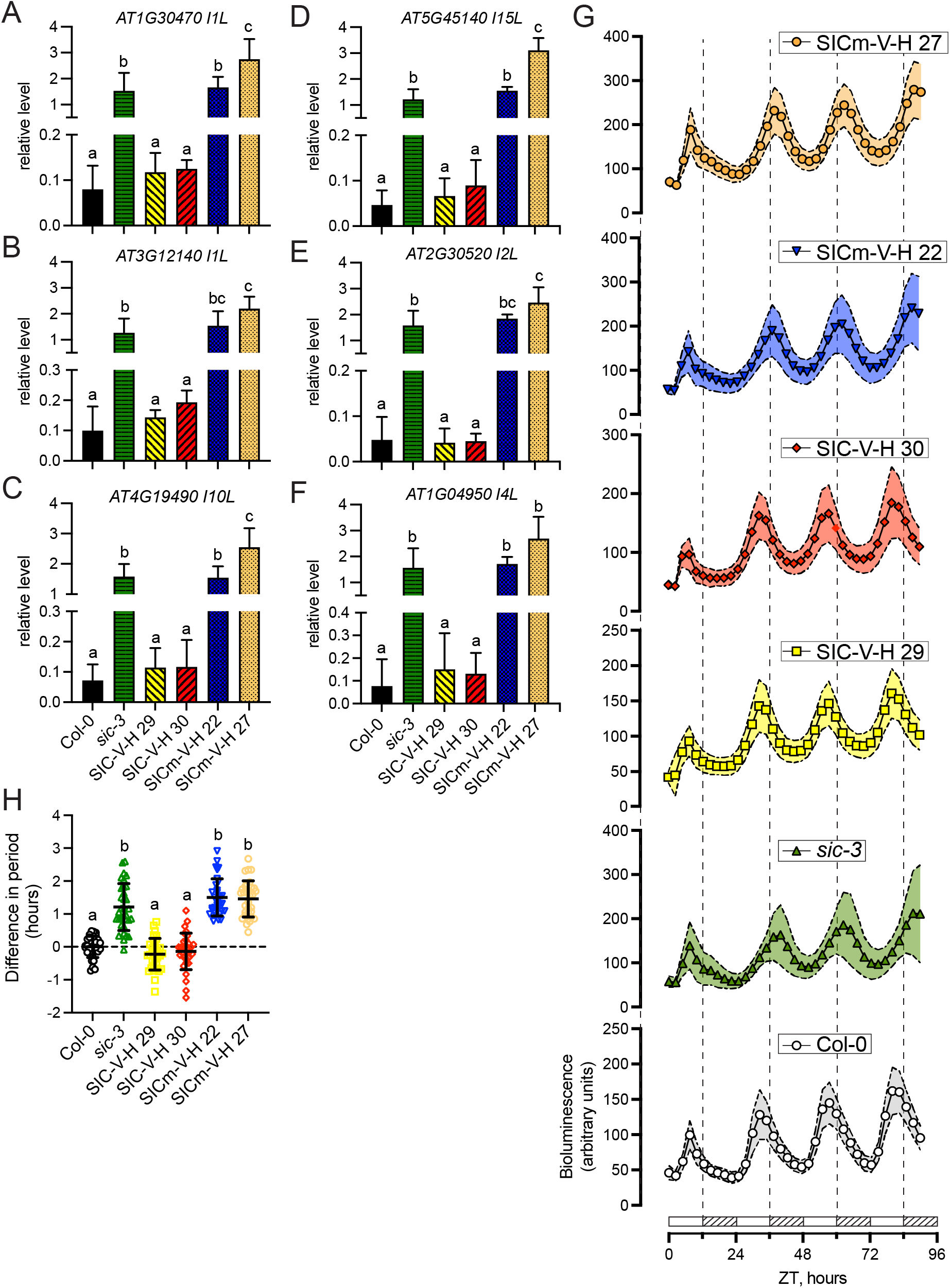
The SIC MPLKIP motif is needed for activity *in vivo*. A-F) Quantification of intron lariats A) *AT1G30470 I1L*, B) *AT3G12140 I1L* C), *AT4G19490 I10L*, D) *AT5G45140 I15L*, D) *AT2G30520 I2L*, and F) *AT1G04950 I5L* by qPCR from RNase R treated samples of Col-0 (black solid bar), *sic-3* (green horizontal hatched bar), *SIC-V-H 29* (yellow left crosshatched bar), *SIC-V-H 30* (red right crosshatched bar), *SICm-V-H 22* (blue checkered bar) and *SICm-V-H 27* (orange checkered bar). The error bars are standard deviation of 3-4 independent biological replicates. G) Circadian rhythms of the *PRR7:LUC+* reporter in Col-0 (white circles), *sic-3* (green triangles), *SIC-V-H 29* (yellow squares;), *SIC-V-H 30* (red diamonds), *SICm-V-H 22* (blue inverted triangles) and *SICm-V-H 27* (orange circles) under constant white light and 22°C. Bioluminescence of seedlings at the indicated number of hours after transfer to constant conditions (zeitgeber time, ZT). Line and symbols indicate the mean of all seedlings of that genotype and shaded regions indicate standard deviation. White bars and hatched bars on the x-axis indicate subjective day and night, respectively. Vertical dotted lines mark the onset of subjective night. H) Difference in estimated period from the mean of Col-0 (dashed horizontal line). Symbols and colors are the same as in (G). Mean period indicated by dashed line and error bars are standard deviation. Data are from 2 independent biological replicates and n = 32. Period estimates are from individuals with relative amplitude error values <0.6. Means sharing a common letter are not significantly different by one-way ANOVA with Tukey’s multiple comparisons test at p <0.05 level of significance.

## DISCUSSION

Here we demonstrate intron lariats accumulate to high levels in *sic* mutants, a phenotype like *dbr1* mutants. Indeed, *sic* and *dbr1* mutants harbor many of the same stable intron lariats. Since DBR1 is the sole intron lariat debranching enzyme in Arabidopsis (Wang et al., 2004), we conclude SIC is required for debranching of intron lariats by DBR1. In addition, the *sic-1* and *sic-3* mutants incur physiological phenotypes like *dbr1-3*, including inhibited root growth, diminished rosette size, and a slower, less precise circadian clock. Thus, the same spectrum of physiological phenotypes is common to these mutants that amass high levels of intron lariats. Finally, SIC directly interacts with DBR1 and this interaction occurs through the evolutionary conserved MPLKIP domain in SIC. A mutant SIC protein without the MPLKIP motif did not complement intron lariat over production or circadian clock dysfunction in *sic-3*.

Combining the *sic* alleles with the *dbr1-3* allele revealed a complex genetic interaction, interpretation of which was complicated by the hypomorphic nature of each allele. The homozygous *sic-1 dbr1-3* allele could not be made, indicating a potential negative synergistic interaction between these two alleles that produces an embryo or seedling lethal phenotype. This apparent synthetic lethal phenotype may occur for the same reasons the null *dbr1-1* allele is embryo lethal (Wang et al., 2004). The *sic-3 dbr1-3* mutant was readily recovered from crosses, most likely because the *sic-3* allele is weaker than *sic-1* (Marshall et al., 2016). *sic-3* is potentially epistatic to *dbr1-3*, since most of the phenotypes tested in *sic-3 dbr1-3* were as severe as the *sic-3* mutant. The exception was a reduction in circadian clock precision and accuracy in *sic-3 dbr1-3* mutant that was substantially greater than the individual mutants. Thus, *sic-3 dbr1-3* appears to alter the circadian system in ways unique from *sic-3*, *sic-1,* and *dbr1-3*. The nature of these changes is not clear but may involve alternative splicing of key circadian clock transcripts (Marshall et al., 2016) and/or consequences of changes in the accumulation of small RNAs like miRNAs (Zhan et al., 2012) or possibly other types of regulatory RNAs.

The discovery of intron lariat accumulation in *sic* mutants identifies the potential root cause for the pri-miRNA and pre-mRNA processing defects identified in *sic-1* and *sic-3*. Amassing of intron lariats in *sic-1* accounts for the discovery of undegraded introns in *sic-1* (Zhan et al., 2012). Furthermore, inhibition of pri-miRNA processing and the consequent reduction in miRNA production in *sic-1* (Zhan et al., 2012) can now be explained by the same mechanism that impairs miRNA production in *dbr1-2* (Li et al., 2016). High levels of intron lariats in *dbr1-2* inhibit dicer complex-mediated processing of pri-miRNAs into miRNAs. The cause of dicer complex inhibition is a “molecular sponge” effect whereby intron lariats out compete pri-miRNAs for binding to the core dicer complex components DCL1 and HYL1. It is reasonable to conclude that the high levels of intron lariats in *sic-1*, and likely in *sic-3*, similarly inhibit DCL1/HYL1 activity. Consistent with this idea, *sic-1* and *dbr1-2* share reductions in many of the same miRNAs, including *miR159*, *miR165*, *miR168*, *miR172* and *miR173* (Li et al., 2016; Zhan et al., 2012).

Both *sic* and *dbr1* mutants also interfere with pre-mRNA processing as indicated by alternative splicing of transcripts. Previous work has found *sic-1* and *sic-3* exhibit changes in mRNA splice variant abundance and diversity (Marshall et al., 2016; Zhan et al., 2012). DBR1 activity is linked to pre-RNA processing in both yeast and human cells. The *Schizosaccharomyces pombe* null *dbr1* mutant accumulates intron lariats and exhibits reduced splicing efficiency for many introns (Bitton et al., 2014). Human DBR1 activity is critical for efficient pre-mRNA processing because of the role it plays in recycling of spliceosome components associated with intron lariats at the completion of splicing. When DBR1 activity is low, the Prp19C/NTC and U5 snRNP subunits become trapped in post-splicing complexes with intron lariats instead of being reused in succeeding active spliceosome complexes associated with pre-mRNAs (Han et al., 2017). The consequence is reduced availability of core spliceosome components like the Prp19C/NTC and U5 snRNP, which leads to skipping of exons with weak splice sites and, ultimately, transcript splice variants (Han et al., 2017). Such defects in RNA processing explain, in part, why inappropriate regulation of *DBR1* is associated with oncogenesis in several types of human cancer cells (Han et al., 2017). We propose stable intron lariats in Arabidopsis *sic* and *dbr1* mutants interfere with pre-mRNA processing and this leads to alternative splicing of transcripts.

SIC is clearly important for lariat debranching by DBR1, but its molecular function remains to be identified. Because SIC directly interacts with DBR1, it likely is a key co-factor of DBR1; however, SIC may instead provide DBR1 with appropriate intron lariat substrates during transcript splicing. A model for SIC function is one where intron lariat processing is mediated by a multiprotein complex in which SIC directly associates with DBR1 as recently described for the human TTDN1 and DBR1 proteins. TTDN1 recruits human DBR1 to the intron-binding complex which contains core Prp19C/NTC subunits including the Aquarius helicase (AQR), XAB2/ SYF1, and ISY1 (Townley et al., 2023). Similarly, a previous SIC TAP-MS experiment that isolated DBR1 yielded many peptides from the AQUARIUS, ISY1, and XAB2 (Karampelias et al., 2016). Human DBR1 interacts with human XAB2 in the intron large complex (Masaki et al., 2015), which is a post-splicing complex related to the intron lariat spliceosome complex. In yeast, the intron lariat spliceosome complex consists of an intron lariat associated with several multiprotein assemblies including the Prp19C/NTC (Wan et al., 2017). Thus, SIC and Arabidopsis DBR1 are likely to participate in the plant equivalent of the post-splicing intron lariat spliceosome complex.

Interestingly, disassembly of the intron lariat spliceosome complex is necessary for human and yeast DBR1 to access a bound intron lariat. *In vitro* studies with purified human intron lariat spliceosome complexes demonstrate human DBR1 does not efficiently debranch intron lariats bound by Prp19C/NTC, U2 snRNP and U5 snRNP subunits (Yoshimoto et al., 2009). But DBR1 efficiently debranches intron lariats in post-splicing RNA-protein complexes lacking these components. Arrested splicing complexes accumulate in a *Saccharomyces cerevisiae* mutant lacking Prp43, a key post-splicing complex disassembly factor, and the intron lariats within these complexes are resistant to DBR1 *in vitro* (Martin et al., 2002). A possibility is SIC promotes DBR1 activity by providing it access to intron lariats in a post-splicing complex like the intron lariat spliceosome complex. A comparable role is proposed for human TTDN1 (Townley et al., 2023).

In summary, these findings indicate SIC is needed for prompt intron lariat removal at the completion of splicing. Since persistent intron lariats interfere with pri-miRNA and pre-mRNA processing, SIC also is indirectly required for these activities. Therefore, SIC is a critical factor for maintenance of RNA homeostasis in Arabidopsis. This work, together with discovery of human TTDN1 protein function, indicates a broadly conserved role for MPLKIP motif containing proteins in the mechanisms that dictate the fate of intron lariats, and possibly the fate of post splicing spliceosome complexes, across plants and animals. Consequently, appreciation of SIC function will reveal important yet understudied facets of spliceosome function in plants and animals.

## METHODS

### Plant material, growth, and phenotype analysis

All plant material was the *Arabidopsis thaliana* Col-0 accession with the *PRR7:LUC+* bioluminescent reporter construct (Salome & McClung, 2005). The *sic-1* allele was from Zhan et al. (Zhan et al., 2012) and the *sic-3* allele was from Marshall et al. (Marshall et al., 2016). The *dbr1-3* allele was from the SALK T-DNA insertion collection (Alonso et al., 2003), and corresponds to line SALK_041038 (SFigure 4A) obtained from the Arabidopsis Biological Resource Center (Columbus, OH). The *PRR7:LUC+* reporter was introduced into *dbr1-3* by crossing to Col-0 carrying the reporter. To generate *sic dbr1-3* double mutant combinations, emasculated *sic* flowers were pollinated with anthers from homozygous *dbr1-3* mutant flowers. The presence of *sic-1* and *sic-3* alleles was detected by restriction digestion of PCR products amplified from genomic DNA with primers sic1dCAPXcmIF and sic1dCAPXcmIR (STable I). The *sic-1* mutation blocks digestion by enzyme *XcmI* and *sic-3* blocks digestion by enzyme *BstNI*. The presence of the *dbr1-3* allele was detected by PCR amplification from genomic DNA with primers LBb1.3 (Alonso et al., 2003) and SALK_041038R (STable I). The absence of the *dbr1-3* allele was detected with primers SALK_041038F and SALK_041038R.

Plants were started on MS plates composed of 1X Murashige and Skoog basal salt medium (Sigma-Aldrich, St. Louis, MO) at pH 5.7-5.9 with 0.8% Type I micropropagation agar (Spectrum Laboratory Products, Inc., New Brunswick, NJ). Plates were 100 x15 mm square plates (VWR Scientific Products, Randor, PA). Surface sterilized seeds were prepared by a 10-minute treatment with an aqueous solution of 30% bleach and 0.01% SDS followed by extensive washing with sterile Milli-Q water (EMD Millipore, Hayward, CA). Seeds were then sown onto MS plates and stratified by wrapping the plates in aluminum foil and placing them at 4°C for 3 days.

For root growth analysis, seeds of indicated genotypes were sown along the top of the MS plates. Following stratification as above, plates were transferred to constant light and 22°C (LL|22°C), oriented vertically with seeds at the top. After 3 days under LL|22°C, conditions were changed to 12 hours light/12 hours darkness and 22°C (LD|22°C). Cool white fluorescent bulbs provided light at 50 μmol photons m^-2^s^-1^. When seedlings reached 7-days-old, the plates were digitally photographed. Root length was determined with ImageJ (Abramoff et al., 2004) by measuring the length of a line drawn along the root. The value for each seedling was normalized to the average for Col-0.

For rosette diameter measurements, plates with seeds of indicated genotypes stratified as above were transferred to LL|22°C and kept there for 3 days, then conditions were changed to LD|22°C. Once seedlings reached 7-8 days old, they were transplanted to 27.8 x 54.5 x 6.2 cm trays with 4 x 6 inserts filled with Sunshine soil (The Scotts Company, Marysville, OH) wetted with tap water. Plants were grown to 20 days old under 16 hours light/8 hours darkness and 22°C in the Plant Gene Expression Center greenhouse in Albany, CA, between June 2020 and August 2020. Supplemental lighting was provided by LED grow lights (Lumigrow, Emeryville, CA), with intensity set at 3. Plants were digitally photographed, and rosette diameter was measured with Image J (Abramoff et al., 2004). Rosette diameter was calculated from the circumference of a circle drawn to the edge of each plant rosette. Measurements were normalized to the average for Col-0.

### RT-PCR and qPCR for lariat detection, quantification, and sequencing

Plants were grown to 8 days old as described above. At 2 hours after dawn (zeitgeber time 2, ZT2), whole plants were collected and flash-frozen in liquid nitrogen. Three biological replicates were collected for each genotype at each time point. After the tissue was ground under liquid nitrogen, total RNA was extracted with Plant RNA Reagent (ThermoFisher Scientific, Waltham, MA) according to the manufacturer’s recommendations. For RNase R digestion, a 4-microgram total RNA sample was prepared in 1X RNase R buffer and split into equal portions, one 2 microgram portion received 5 units of RNase R (Lucigen, Middleton, WI) and the other an equal volume of nuclease-free water (Thermo Fisher Scientific, Waltham, MA). Samples were incubated at 37°C for 30 minutes. Each 2-microgram sample of RNA was treated with dsDNase (ThermoFisher Scientific, Waltham, MA) to remove contaminating genomic DNA.

Preparation of cDNA from the RNA employed the Maxima H Minus First Strand cDNA Synthesis Kit (ThermoFisher Scientific, Waltham, MA) with random primer according to the manufacturer’s recommendations and the final products diluted with 2 volumes of Milli-Q water (EMD Millipore, Hayward, CA). This final cDNA sample served as a template for RT-PCR and qPCR with the primers listed in STable 1.

qPCR involved two technical replicate reactions composed and performed as described previously (Bendix et al., 2013). C_q_ values were calculated with the regression function for each primer set in the Bio-Rad CFX Manager Software (Bio-Rad, Hercules, CA). Values of relative transcript levels were calculated as 2∧(C_q_^normalizer^-C_q_^experimental^). Where C_q_^normalizer^ is the geometric mean of the C_q_ values for the *IPP2* and *PP2A* primer sets for cDNA made from total RNA not treated with RNase R. Individual relative transcript level values were normalized to the average of values from all genotypes and treatments (RNase R treated and untreated). Sanger sequencing of intron lariat qPCR products employed the forward primer of the pair.

### Circadian clock analysis

Circadian clock behavior under constant white light and temperature (22°C) conditions (LL|22°C) was determined by time-lapse imaging of bioluminescence from the *PRR7:LUC+* reporter (Salome & McClung, 2005). Seedlings were grown to 8 days old on MS plates under LD|22°C conditions, after stratification and 3 days under LL|22°C as described above. The day before imaging, each plate was sprayed with 1 milliliter of an aqueous solution of 5 mM firefly luciferin (Gold Biotechnology, St Louis, MO) and 0.01% (vol/vol) Triton X-100 (Sigma Aldrich, St. Louis, MO). Plates were transferred to LL|22°C (50 μmol photons m^-2^s^-1^ from halogen bulbs) at ZT0 and imaged every 2.5 hours with an ORCAII camera (Hamamatsu Photonics, Hamamatsu City, Japan). Bioluminescence of individual seedlings was collected and extracted from images with MetaMorph software (Molecular Devices, San Jose, CA). Fast Fourier Transform-Nonlinear Least Squares (FFT-NLLS) analysis within the Biological Rhythms Analysis Software System 3.0 (BRASS) (Locke et al., 2005; Southern & Millar, 2005) was used to calculate the estimated circadian period and relative amplitude error from the experimental bioluminescence time-series data (Plautz et al., 1997). Four independent biological replicates were performed with 16-40 individuals of each genotype included in each replicate. Period estimates were kept only for individuals with RAE values <0.6.

### Site-directed mutagenesis of SIC MPLKIP motif

Site-directed mutagenesis to produce the SICm variant with the core MPLKIP sequence S M A E D P W converted to A A A A A A A employed Splicing by overhang-extension PCR (SOE PCR) (Higuchi et al., 1988; Ho et al., 1989). Primers “SIC_MPLKIP_mut_F” and “SIC_MPLKIP_mut_R” have a 21-nucleotide region of complementarity that substitutes the codons encoding the MPLKIP motif (5’-TCT ATG GCT GAA GAT CCA TGG-3’) with codons for 7 sequential alanine residues (5’-GCT GCA GCT GCT GCA GCG GCA-3’). The final 1,869 base pair SICp:SICm SOE PCR product, which includes the genomic sequence beginning at the end of the gene just upstream of SIC to the end of the SIC coding sequence (without the stop codon) was made with Q5 High-Fidelity DNA Polymerase (New England Biolabs) in three steps. In step 1, fragment 1 and fragment 2 were separately amplified from Col-0 genomic DNA according to the manufacturer’s recommendations. Fragment 1, amplified with primers AT4G24500-124F and SIC_MPLKIP_mut_R, encompasses 392 nucleotides upstream to 1,273 nucleotides downstream from the *SIC* start codon. Fragment 2, amplified with primers SIC_MPLKIP_mut_F and WARP2cdsnostopR, encompasses 1,253 to 1,477 nucleotides downstream of the *SIC* start codon. Each fragment was agarose gel purified with the Qiagen QIAquick Gel Extraction kit. In step 2, 1 ng each of purified fragments 1 and 2 were used as primer and template in a 25 µL PCR reaction with an annealing temperature of 70°C and 10 cycles of amplification. In step 3, 25 µL of the PCR reaction from step 2 was mixed with 25 µL of a standard PCR reaction containing primers AT4G24500-124F and WARP2cdsnostopR with an annealing temperature of 60°C and 35 cycles of amplification. The wild-type version of the same SIC sequence was amplified from Col-0 genomic DNA with primers AT4G24500-124F and WARP2cdsnostopR. These final PCR fragments were gel purified and cloned into the pENTR/D-TOPO vector (ThermoFisher Scientific, www.thermofisher.com) vector as described above, then sequenced to confirm the site-directed mutagenesis. In subsequent cloning, the SICm mutation was identified by restriction digest with the enzyme *PvuII* (New England Biolabs, www.neb.com), since the SICm mutation introduces this site into the *SIC* sequence.

### Yeast two-hybrid

The coding sequences of *SIC* (AT4G24500) and *DBR1* (AT4G31770), including the stop codon sequence, were amplified out of cDNA from *Arabidopsis* Col-0 or an *SICp:SICm-Venus-HA* transgenic line (see below) by PCR with Q5 High Fidelity Polymerase (New England Biolabs, www.neb.com). Primers “WARP2cdsGATEF” and “WARP2cdsR” were for *SIC* and primers “DBR1_CDS_nostp_CACC_F” and “DBR1_CDS_stp_R” were for *DBR1* (Table S1). PCR products were cloned into the pENTR/D-TOPO vector (ThermoFisher Scientific, www.thermofisher.com) and transformed into One Shot TOP10 Chemically Competent Escherichia coli (Thermo Fisher Scientific, www.thermofisher.com). Sequences were confirmed by Sanger sequencing. The *DBR1* and *SIC* coding sequences were subcloned into bait vector pGBKT7-GW and the prey plasmid pGADT7-GW (Lu et al., 2010), respectively, with LR Clonase II (ThermoFisher Scientific, www.thermofisher.com). Negative control LAM pGBKT7 and LAM pGADT7 were from Clontech (Takara Bio USA, Inc.,www.takarabio.com). Bait and prey plasmids were transformed into Y187 and Y2H Gold cells according to the manufacturer’s recommendations (Takara Bio USA, www.takarabio.com). Bait and prey combinations were made by mating Y187 bait and Y2H Gold prey strains according to the manufacturer’s recommendations (Takara Bio USA, www.takarabio.com).

For interaction tests, three individual transformants for each bait-prey combination were grown at 30°C in liquid Synthetic Dropout (SD; Takara Bio USA, www.takarabio.com) media lacking amino acids Trp (W) and Leu (L) (SD-WL) to an absorbance at 595 nm = 1.0, which corresponds to 3x10^4^ cells/μL. A total of 1x10^8^ cells were harvested by centrifugation and resuspended to 1x10^6^ cells/μL with sterile TE buffer (10 mM Tris-HCl, 1 mM EDTA, pH 7.5.). Serial dilutions of this 1x10^6^ cells/μL culture were made in TE buffer to achieve cell densities of 1x10^5^, 1x10^4^, and 1x10^3^ cells/μL. These cultures were spotted in 3 μL onto control SD-WL plates and test SD plates lacking amino acids W, L, and adenine (A) (SD-WTA). After drying, plates were sealed with Micropore Paper Tape (3M, www.3m.com) and placed in a 30**°**C dry air incubator. Digital images of plates were taken after 3, 4, and 5 days to monitor yeast growth.

### *In vitro* co-immuno purification

The mVenus coding sequence was amplified from vector pK-mVenus-HHC (Huang et al., 2016). To construct pT7CFE1-cHis-(ThermoFisher Scientific, www.thermofisher.com) based vectors for *in vitro* protein expression, primers carrying necessary restriction enzyme cut sites and sequences encoding either the HA epitope (YPYDVPDYA) or the cMyc epitope (EEQKLISEEDL) were employed to amplify the *SIC*, *DBR1*, and *mVenus* coding sequences. by PCR with Q5 High-Fidelity DNA Polymerase (New England Biolabs, www.neb.com) polymerase. The desired PCR products were purified from agarose purified and cloned into the pENTR/D-TOPO vector (ThermoFisher Scientific, www.thermofisher.com) and transformed into One Shot TOP10 Chemically Competent Escherichia coli (ThermoFisher Scientific, www.thermofisher.com). The restriction enzyme-epitope-coding sequence fragment was excised by restriction enzyme digestion, isolated by agarose gel electrophoresis, cut from the gel, and purified with the QIAquick Gel Extraction Kit (Qiagen, www.qiagen.com). Fragments were ligated with T4 DNA ligase (New England Biolabs, www.neb.com) into linearized pT7CFE1-cHis digested with complementary restriction enzymes and transformed into 5-alpha Competent E. coli cells (New England Biolabs, www.neb.com). Sequences were confirmed by Sanger sequencing.

Proteins for CoIP were produced with the TnT Quick Coupled Transcription/Translation System (Promega, www.promega.com) according to the manufacturer’s recommendations. The HA-SIC and HA-SICm bait protein product were incubated on a rotator with 25 μL of Pierce Anti-HA Magnetic Beads (ThermoFisher Scientific, www.thermofisher.com), equilibrated in cold PBS buffer (10 mM Na_2_HPO_4_, 1.8 mM KH_2_PO_4_, 137 mM NaCl, 2.7 mM KCl, 1X Roche cOmplete EDTA-free protease inhibitor cocktail (ThermoFisher Scientific, www.thermofisher.com), and 0.1% NP-40), for 40 minutes at room temperature then washed three times with 10 volumes of cold PBS buffer. The cMyc-DBR1 and cMyc-mVenus prey proteins were incubated with the prepared beads on a rotator for 1 hour at 4C. Beads were washed three times with 10 volumes of cold PBS buffer. Proteins on the beads were released by incubation at 95C with 100 μL of 1X nonreducing sample buffer (60 mM Tris-HCL, pH 6.8, 1% SDS, 10% glycerol and 0.02% bromphenol blue). The supernatant was removed and equilibrated to 5 mM DTT. Proteins detected with western blot analysis as described in Supplemental Methods.

### Construction of *SICp:SIC/SICm-Venus-HA* transgenic lines

The *SICp:SIC* and *SICp:SICm* sequences in pENTR/D-TOPO (described above) were subcloned into binary vector pK-mVenus-HHC (Huang et al., 2016) with LR Clonase II (ThermoFisher Scientific, www.thermofisher.com).The *SICp:SIC-mVenus-HA* and *SICp:SICm-mVenus-HA* vectors were transformed into *sic-3* plants by floral dip (Clough & Bent, 1998). T_1_ transformants were selected for by growth of surface sterilized seeds on MS plates supplemented with 16 µg/ml phosphinothricin (PhytoTech Labs, www.phytotechlab.com) as described above. After 10 days, resistant seedlings were transferred to soil. The presence of the transgene was confirmed by PCR genotyping with primers sic1dCAPXcmIF and EYFPQR. Digestion of this PCR product with *PvuII-HF* (New England Biolabs, www.neb.com) distinguished between SICm (*PvuII* sensitive) and SIC (*PvuII* resistant). The T_2_-T_5_ generations were grown out to identify homozygous lines and characterize protein expression by western block and fluorescent confocal microscopy.

## ACKNOWLEDGMENTS

We thank Stacey Harmer and Marcelo Yanovsky for valuable discussions and feedback. Ariel Chen provided technical assistance in identification of the *sic-3 dbr1-3* mutant and analysis of *sic-1* X *dbr1-3* crosses. Parkesh Suseendran and Sienna Weinstein assisted in characterization of transgenic lines. Thank you to PGEC greenhouse manager, Lia Poasa, and her staff, Shawna Kelly, Chris Tucker, and Emilio Corona for excellent care of plants.

## AUTHOR CONTRIBUTIONS

E.K., C.M.M. and F.G.H. conceived and designed the experiments; C.M.M., E.K., and F.G.H performed the experiments. M.D.C.M and F.G.H. performed data analysis. A.L.N. provided computational resources. F.G.H. wrote the first draft of the manuscript. E.K., C.M.M., M.D.C.M, A.L.N. and F.G.H. edited the text and approved the final manuscript.

## SUPPLEMENTARY INFORMATION

### SUPPLEMENTARY TABLES

**STable 1. Primers used in this study.**

**STable 2. Summary statistics of RNAseq data sets.**

**STable 3. Features with differential usage in *sic-3*.**

**STable 4. Pearson correlation coefficients between K-means cluster centroids generated with the indicated K values.**

**STable 5. K-means cluster membership from ShapeShifter analysis.**

**STable 6. ASpli designation of RNase R resistant intron sequences from *dbr1-2*.**

**STable 7. Features with differential usage in *dbr1-2*.**

### SUPPLEMENTARY FIGURES

**SFigure 1. RNAseq profiling reveals accumulation of intron sequences in sic-3.**

**SFigure 2. ShapeShifter analysis pipeline identifies intron sequences with intron lariat attributes.**

**SFigure 3. Pretreatment with RNase R effectively removes mRNA from total RNA.**

**SFigure 4. Sequences of RT-PCR products amplified from intron lariats in sic-3.**

**SFigure 5. Characterization of the dbr1-3 allele.**

**SFigure 6. ASpli analysis of RNAseq profiling of dbr1-2 and Col-0.**

**SFigure 7. Alignment of MPLKIP motifs from plants, fungi, and animals.**

**SFigure 9. Characterization of SICp:SIC-mVenus-HA/sic-3 and SICp:SICm-mVenus-HA/sic-3 transgenic lines.**

### SUPPLEMENTARY METHODS

**Growth conditions and RNAseq profiling of *sic-3* and Col-0**

**Read alignment and ASpli analysis to identify differential usage splice variants in *sic-3* and *dbr1-2*.**

**ShapeShifter analysis pipeline to identify intron lariat profiles in *sic-3***

**Analysis of epitope-tagged protein accumulation in transgenic lines**

**Western blot**

**Confocal microscopy**

**Statistical analysis**

### SUPPLEMENTARY REFERENCES

