## Supplementary Text, Figures for "Lariat debranching by RNA DEBRANCHING ENZYME 1 depends on SICKLE in *Arabidopsis thaliana*"

### SUPPLEMENTARY METHODS

#### Growth conditions and RNAseq profiling of *sic-3* and Col-0

For the RNAseq profiling comparison of *sic-3* to Col-0, seeds on MS plates were stratified at 4°C for 3 days, moved to LL|22°C for 3 days, moved LD|22°C for 7 additional days, and on the last day these 10-day-old seedlings were transferred to LL|16°C at 12 hours after dawn (ZT 12). After 8 hours of acclimatization to the new temperature, tissue collection began and occurred every 4 hours for the next 20 hours (samples at ZT 20, ZT 24, ZT 28, ZT 32, ZT 36, and ZT 40) (SFigure 1A). Three biological replicates were collected from three concurrent experiments. Light was provided by cool white fluorescent bulbs at 50 µmol photons $m^{-2} s^{-1}$. Total RNA was extracted with Plant RNA Reagent according to the manufacturer's protocol (Thermo Fisher Scientific, www.thermofisher.com). RNA concentration was measured with the Qubit RNA BR Assay Kit (Thermo Fisher Scientific, www.thermofisher.com) and an RNA pool made for *sic-3* and Col-0 from equal RNA amounts from each time point. poly(A) RNA was purified from 3 micrograms of each RNA pool with the Dynabeads Oligo(dT) kit (Thermo-Fisher Scientific, www.thermofisher.com) and this purification was repeated. RNAseq libraries were prepared from the poly(A) RNA fraction with the ScriptSeq v2 RNAseq library preparation kit (Epicentre, epicentre.com), and the libraries were indexed using ScriptSeq Index PCR primers (Epicentre, epicentre.com) according to the manufacturer's recommendations. Paired-end sequencing with 150 cycles was performed by Illumina HiSeq4000 at the UC Davis DNA Technologies & Expression Analysis Core Laboratory (Davis, CA). The FASTQ files from Illumina sequencing are available as a BioProject at the National Center for Biotechnology Information (NCBI) under accession number PRJNA758710 (Marshall & Harmon, 2021).

**Read alignment and ASpli analysis to identify differential usage of splice variants in *sic-3* and *dbr1-2*.**

The FASTQ files for *sic-3* and matched Col-0 samples were from the RNAseq experiment described above. The RNAseq FASTQ files for *dbr1-2* and matched Col-0 samples were from NCBI BioProject PRJNA291954 and were downloaded from the NCBI Sequence Read Archive (SRA). Files used in ASpli analysis of differential usage splice variants between *dbr1-2* and Col-0 were from RNA samples that were not RNase R treated (SRA accession numbers SRR2145231 and SRR2145235 for Col-0 and SRR2145236 and SRR2145238 for *dbr1-2*.). Files used to generate read pileup images were from RNase R treated RNA samples from *dbr1-2* and Col-0 (SRA accession numbers SRR2145247, SRR2145248 for *dbr1-2* and SRR2145245, SRR2145246 for Col-0). Untrimmed reads in FASTQ files were aligned to the *Arabidopsis thaliana* TAIR10 genome sequence (EnsemblPlants release-35) with HiSat2 version 2.0.5 (Pertea et al., 2016) utilizing default settings with *--rna-strandedness* set to *FR* to produce BAM alignment files. Differential usage transcript splice variants were identified with the R implementation of the ASpli computational pipeline version 2.10.0 (Mancini et al., 2014; Mancini et al., 2021). Only features ≥100 base pairs in size were considered, as done previously for the identification of splice variants in Arabidopsis (Hernando et al., 2015). ASpli estimates differential usage with the statistical method of the

Bioconductor edgeR package (Robinson et al., 2009). ASpli considers only expressed genes and adjusts the counts of features for gene-level expression, thereby removing the contribution of gene expression changes from the differential usage analysis. Features with a $\log_2$ fold change increase >1 and a Benjamini-Hochberg (Benjamini & Hochberg, 1995) corrected false discovery rate value $\leq 0.05$ were considered as having differential usage.

**ShapeShifter analysis pipeline to identify intron lariat profiles in *sic-3***

ShapeShifter analysis was modeled after the method described by Taggart and Fairbrother (Taggart & Fairbrother, 2018). The three *sic-3* RNAseq replicates were analyzed separately with the pipeline outlined in SFigure 2A. Input data were read coverage for the sense strand as a bigWig file, calculated for 10 base pair bins by the bamCoverage tool version 3.4.3 from the HiSat2-generated BAM alignment files described above, and position information for 47,773 introns in a BED file, generated from all introns considered by the ASpli pipeline above. From these data, the computeMatrix tool version 3.4.3 calculated the number of reads aligned to introns in sequential 10 bp bins that began at the predicted 5' end of the intron, and extended to 110 base pairs downstream, with a minimum read depth threshold of 10 (--referencePoint TSS --beforeRegionStartLength 0 --afterRegionStartLength 110 --binSize 10 --minThreshold 10). bamCoverage and computeMatrix are part of the deepTools suite (Ramirez et al., 2016). Read coverage for the bins within intron regions ≥100 bp in length was scaled and centered with the R 'scale' function. The total number of introns passing these filters was 7,949 for replicate 1, 7,047 for replicate 2, and 7,233 for replicate 3. To identify regions with similar read coverage patterns, K-mean clustering of scaled and centered data was performed with the R 'kmeans' function and the cluster cores as centroids with the 'clust.centroid' function. Calculation of sum of squared error, average silhouette width, the Calinski-Harabasz index, gap statistic, and hierarchical clustering in R indicated the optimum number of K-means clusters was between 4 and 7. To select the optimal number of clusters, clustering was done independently with cluster number (K) set to 4, 5, 6, and 7. Analysis of the Pearson correlation coefficient between cluster centroids from each K value indicated K = 5 produced distinct clusters at a significance cutoff of <0.75 (STable 4). These 5 clusters had distinct patterns of read accumulation (Figure 2B) and these patterns appeared for each of the three replicates. Consensus clusters 1 through 5 contain introns that co-occur in all three replicates and in clusters with the same read accumulation profile in all three replicates (STable 5). A total of 2,576 introns were in the consensus clusters (STable 5)

**Analysis of epitope-tagged protein accumulation in transgenic lines**

Seedlings were grown on MS plates under standard conditions. At two weeks, seedlings were harvested at 2 hours after lights on (ZT2) and immediately frozen in liquid nitrogen. Tissue was ground with a mortar and pestle under liquid nitrogen. Whole cell extracts were prepared from ground tissue as described previously (Martínez-García et al., 1999). Three volumes of extraction buffer (100 mM MOPS pH7.6, 100 mM

NaCl, 5% vol/vol SDS, 5 mM EDTA, 5 mM DTT, 10% vol/vol glycerol, and 1X Roche cOmplete EDTA-free Protease Inhibitor Cocktail (Sigma-Aldrich, www.sigmaaldrich.com), heated to 95°C was added to 1 volume of frozen ground tissue, mixed vigorously to wet all tissue, and incubated at 95°C for 5 minutes. Samples were homogenized with a QIAshredder (Qiagen, www.qiagen.com) and the supernatant clarified by centrifugation at 13,000 rpm for 10 minutes at 4°C. Total protein concentration was determined with the Pierce BCA Assay (ThermoFisher Scientific, www.thermofisher.com) according to the manufacturer's recommendations. 60-70 ug of total protein was analyzed by SDS-PAGE and western blot to detect either HA epitope-tagged protein or the tubulin loading control.

**Western blot**

Proteins were separated on a 5% stacking-10% separating (29:1 bis/acrylamide) SDS-PAGE gel. Proteins were transferred to 0.2 micron supported nitrocellulose membrane in transfer buffer (5 mM Tris, 192 mM glycine, 10% methanol) at 4°C. Membranes were blocked with TBSTM (20 mM Tris-HCL, pH 7.5, 150 mM NaCL, 0.1% Tween-20, 5% dry skim milk) at room temperature for at least 1 hour. HA-epitope or cMyc-epitope tagged proteins were detected by probing the membrane with Invitrogen anti-HA antibody (ThermoFisher Scientific, www.thermofisher.com) or Invitrogen anti-cMyc antibody (ThermoFisher Scientific, www.thermofisher.com), respectively, diluted 1:2,500 in TBSTM overnight at 4°C, washing with three changes of TBSTM, and probing with anti-mouse antibody-HRP conjugate (Santa Cruz Biotechnology, www.scbt.com) diluted 1;5,000 in TBSTM at room temperature for 2 hours. After washing with 3 changes of TBST (20 mM Tris-HCL, pH 7.5, 150 mM NaCL, 0.1% Tween-20), membranes were incubated with 1X Amersham ECL Select Western Blotting Detection Reagent (Cytiva, www.cytivalifesciences.com) for 5 minutes. Chemiluminescence was detected with an Odyssey XF molecular imager (Li-Cor, www.licor.com).

**Confocal microscopy**

Detached cotyledons of 7-day-old seedlings grown on MS plates under standard conditions were mounted on microscope slides in water under a coverslip. Samples were imaged with a Leica SP8 confocal laser-scanning microscope with a 40x dry objective. For fluorescence images, the 514 nm argon laser line was used to excite mVenus and florescence observed using the specific emission window of 520–550 nm. The laser power (Argon intensity 25%), gain (1050), zoom (zoom factor 1) and average settings (Format 1024x1024; Speed 200; line average 2; line accuracy 1; frame average 2; frame accuracy 1) were consistent over the same image series to allow fluorescence intensity comparison across samples. Brightfield images were captured with the default settings for Transmitted Light Detection, except the gain was adjusted to maximize contrast. Images were processed using the Leica Application Suite X software package (Leica Microsystems, www.leica-microsystems.com).

**Statistical analysis**

Testing for differences between genotypes for qPCR determined transcript levels, plant measurements, and estimated circadian clock period employed ANOVA with Tukey's multiple comparisons test implemented in Prism 9 (GraphPad, www.graphpad.com) at a significance level of $p < 0.05$. The significance of overlap in membership between datasets was tested with the hypergeometric probability and representation factor as implemented at http://nemates.org/MA/progs/overlap_stats.html. Hypergeometric probability calculated the probability of finding the number of genes shared between two datasets out of the total 47,773 introns considered by the ASpli pipeline. Representation factor = x / expected # of genes, where x = the number of introns in common between two groups and expected number of genes = (n1 x n2) / T, where n1 = number of introns in group 1, n2 = number of introns in group 2, and T = total number of introns (i.e., 47,773). A representation factor > 1 indicates more overlap and representation factor < 1 indicates less overlap than expected for independent groups of genes. Chi-square and Fisher exact testing to evaluate segregation distortion were calculated from contingency tables of expected and observed allele frequencies with chisq.test function in (R Core Team, 2022).

### SUPPLEMENTARY FIGURES

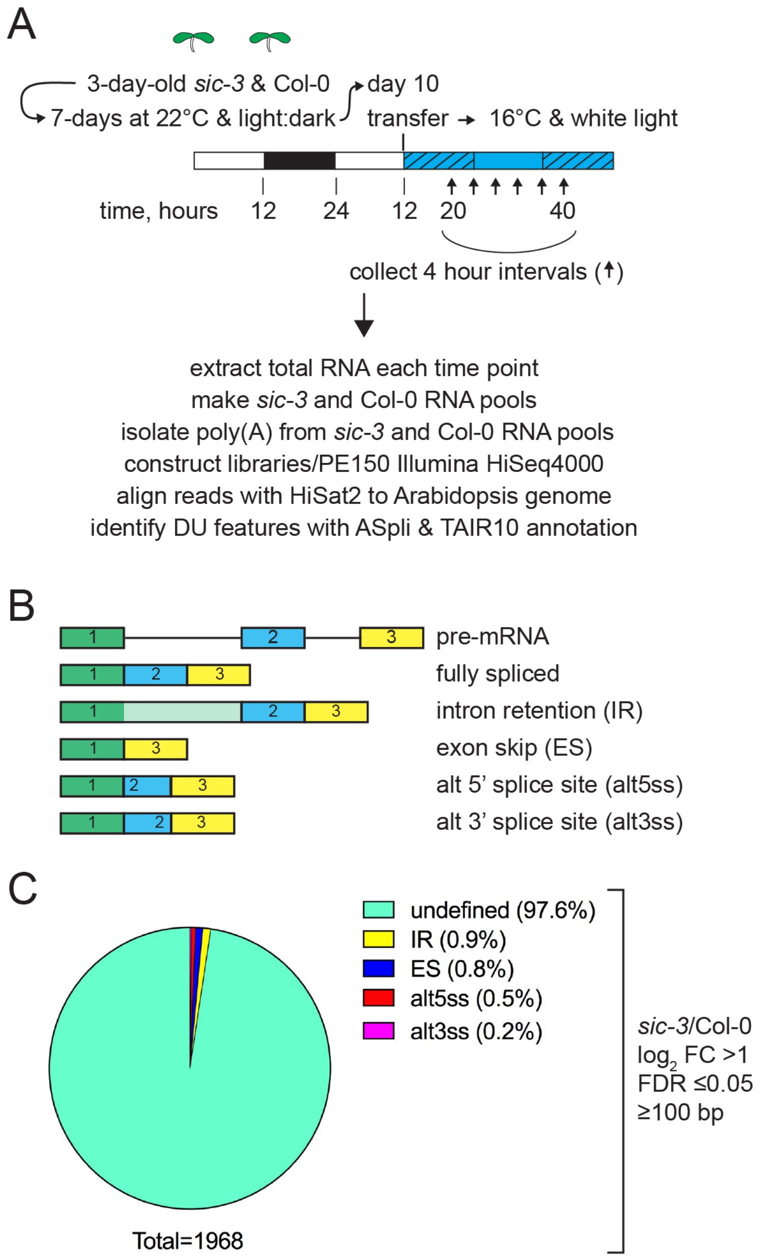


**SFigure 1. RNAseq profiling reveals accumulation of intron sequences in *sic-3*.** A) Growth conditions, sampling plan, and RNAseq profiling protocol for *sic-3* and Col-0 to identify differential usage (DU) of transcript features. White and black bars indicate light and dark conditions, respectively. Hatched bar represents subjective night under constant light conditions. Blue coloring indicates 16°C conditions. B) Definition of splice variant types called by the ASpli pipeline. Colored, numbered boxes are exons and lines are introns. C) Proportion of all *sic-3* differential usage features corresponding to the indicated splice variant types meeting the filtering and statistical criteria shown.

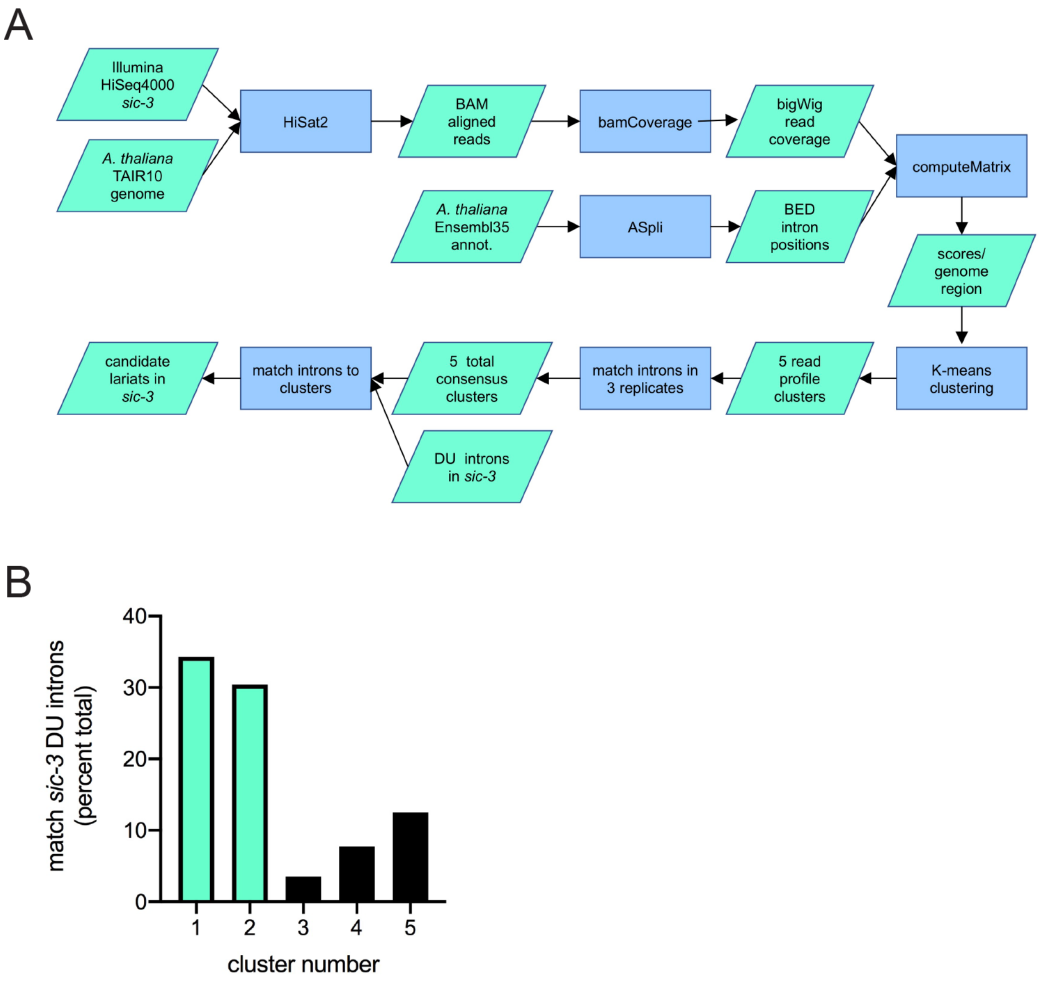


**SFigure 2. ShapeShifter analysis pipeline identifies intron sequences with intron lariat attributes.** A) Flow diagram for ShapeShifter analysis pipeline that identifies intron lariat signatures in RNAseq profiling data. See Supplementary Methods for details. Data set inputs and software tools are shown in green parallelograms and blue rectangles, respectively. B) Proportion of *sic-3* undefined differential usage introns present in ShapeShifter K-means clusters, the majority occur in clusters 1 and 2 (green bars).

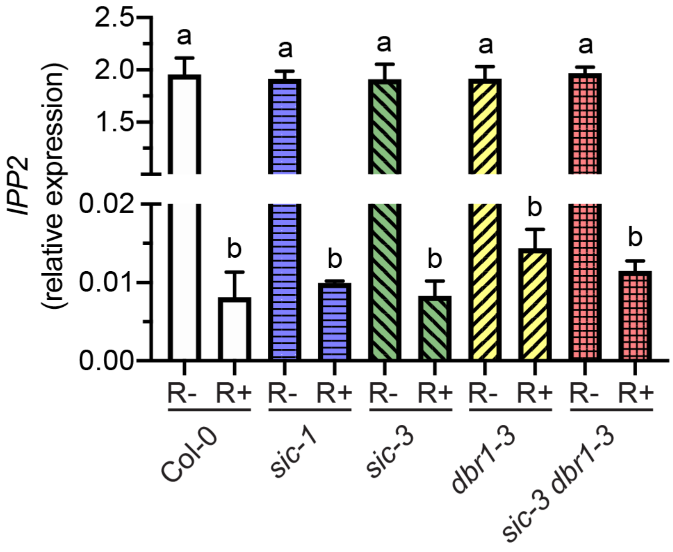


**SFigure 3. Pretreatment with RNase R effectively removes mRNA from total RNA.** Relative expression of *IPP2* transcript determined by qPCR with cDNA prepared with RNA from Col-0 (white solid bar), *sic-1* (blue horizontal hatched bar), *sic-3* (green left crosshatched bar), *dbr1-3* (yellow right crosshatched bar), and *sic-3 dbr1-3* (red checkered bar) without (R-) or with (R+) RNase R pretreatment. Error bars are the standard deviation of three independent biological replicates. Means sharing a common letter are not significantly different by one-way ANOVA with Tukey's multiple comparisons test at $p < 0.05$ level of significance.

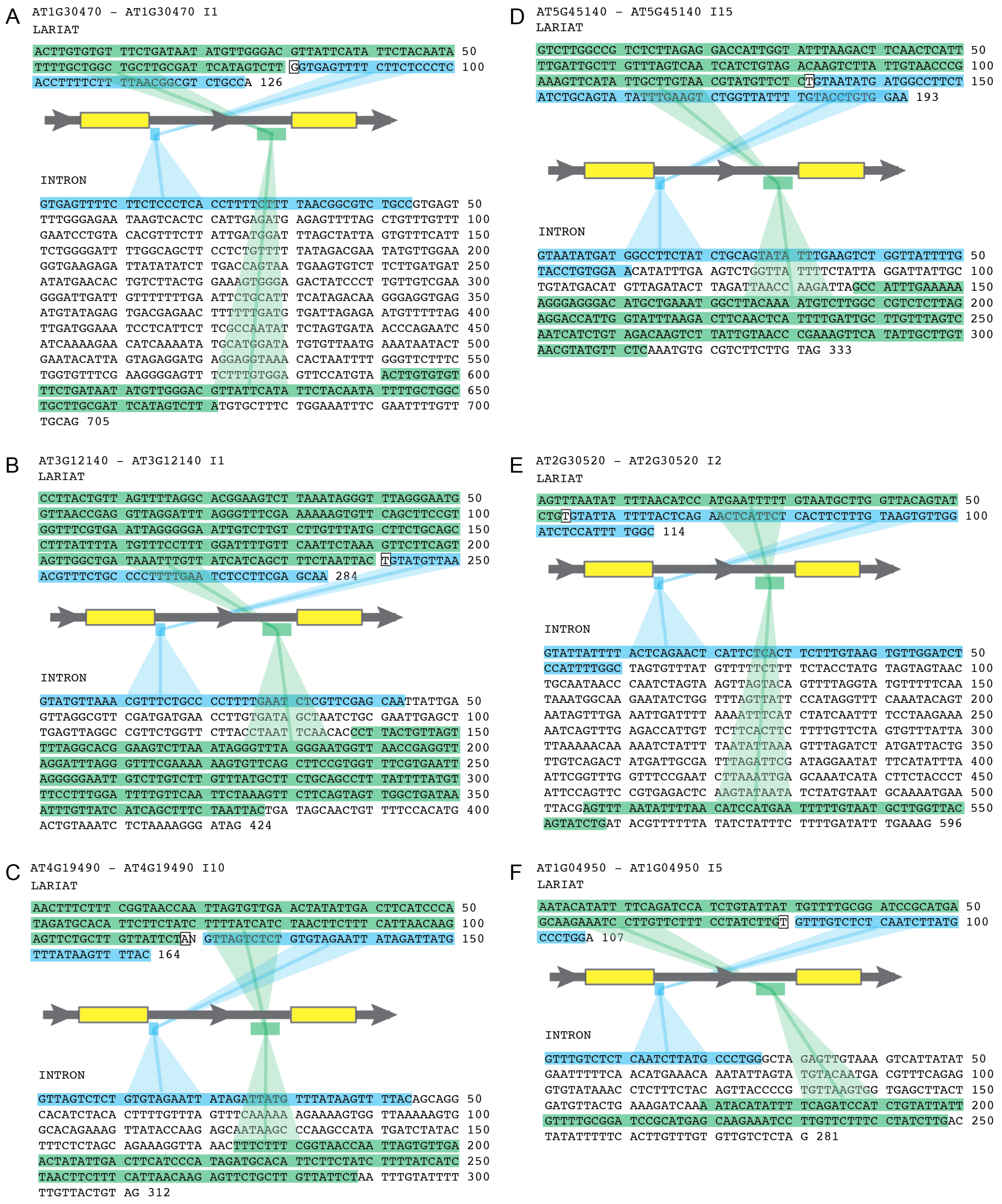


**SFigure 4. Sequences of RT-PCR products amplified from intron lariats in *sic-3*.** Top sequence in each panel is the sequence of the RT-PCR product for A) *AT1G30470 I1L*, B)

*AT3G12140 I1L*, C) *AT4G19490 I10L*, D) *AT5G45140 I15L*, E) *AT2G30520 I2L*, and F) *AT1G04950 I5L* , Bottom sequence is the corresponding intron sequence. Green highlight indicates position of the 5' sequence in the intron lariat that corresponds to the 3' portion of the intron. Blue highlight indicates position of 3' sequence in the intron lariat that corresponds to the 5' end of the intron. Boxed nucleotide in intron lariat is the branchpoint. Cartoon and shaded boxes show relative position of the highlighted sequences in the context of the gene where yellow boxes are exons and grey boxes are introns.

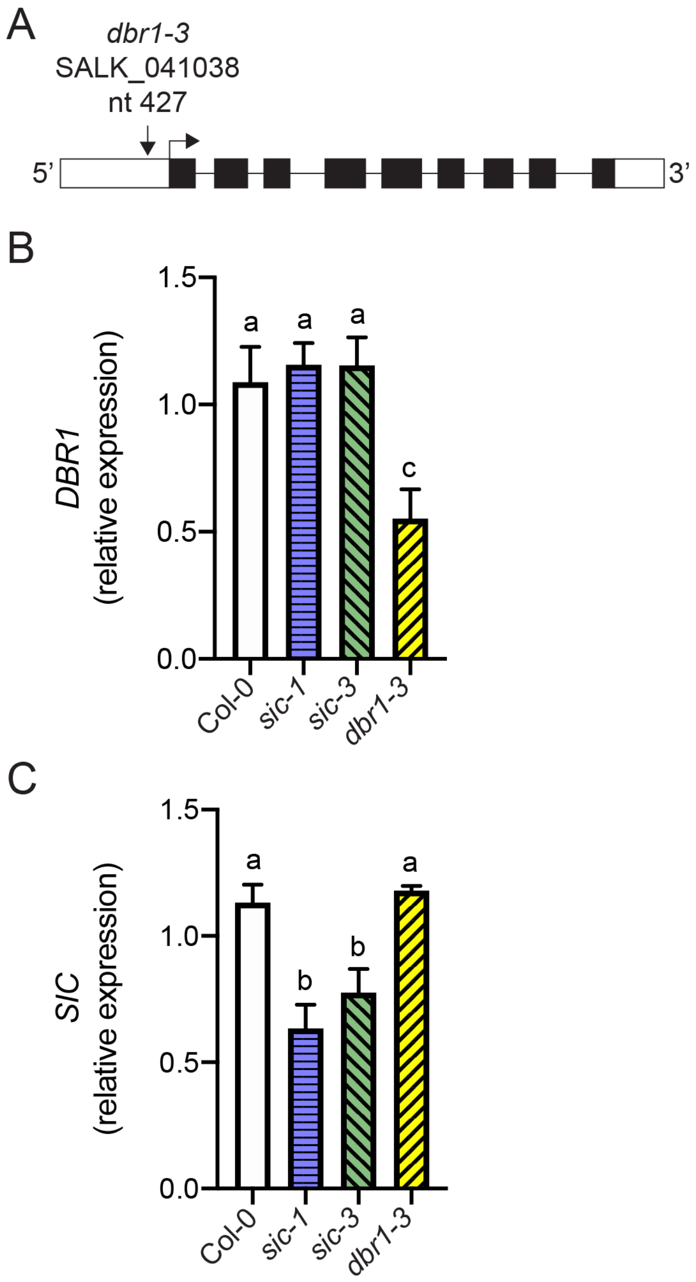


**SFigure 5. Characterization of the *dbr1-3* allele.** A) Schematic of *dbr1-3* allele that bears a T-DNA (SALK_041038) inserted 427 nucleotides downstream of the *DBR1* start (vertical arrow). White boxes are 5' and 3' UTRs and black boxes are exons. Lines are introns. Start codon position indicated by a right-angle arrow pointing to the right. B, C) Relative expression of *DBR1* (B) and *SIC* (C) in Col-0 (white solid bar), *sic-1* (blue horizontal hatched bar), *sic-3* (green left crosshatched bar), and *dbr1-3* (yellow right crosshatched bar) determined by qPCR. Error bars are the standard deviation of three independent biological replicates. Means sharing a common letter are not significantly different by one-way ANOVA with Tukey's multiple comparisons test at $p < 0.05$ level of significance.

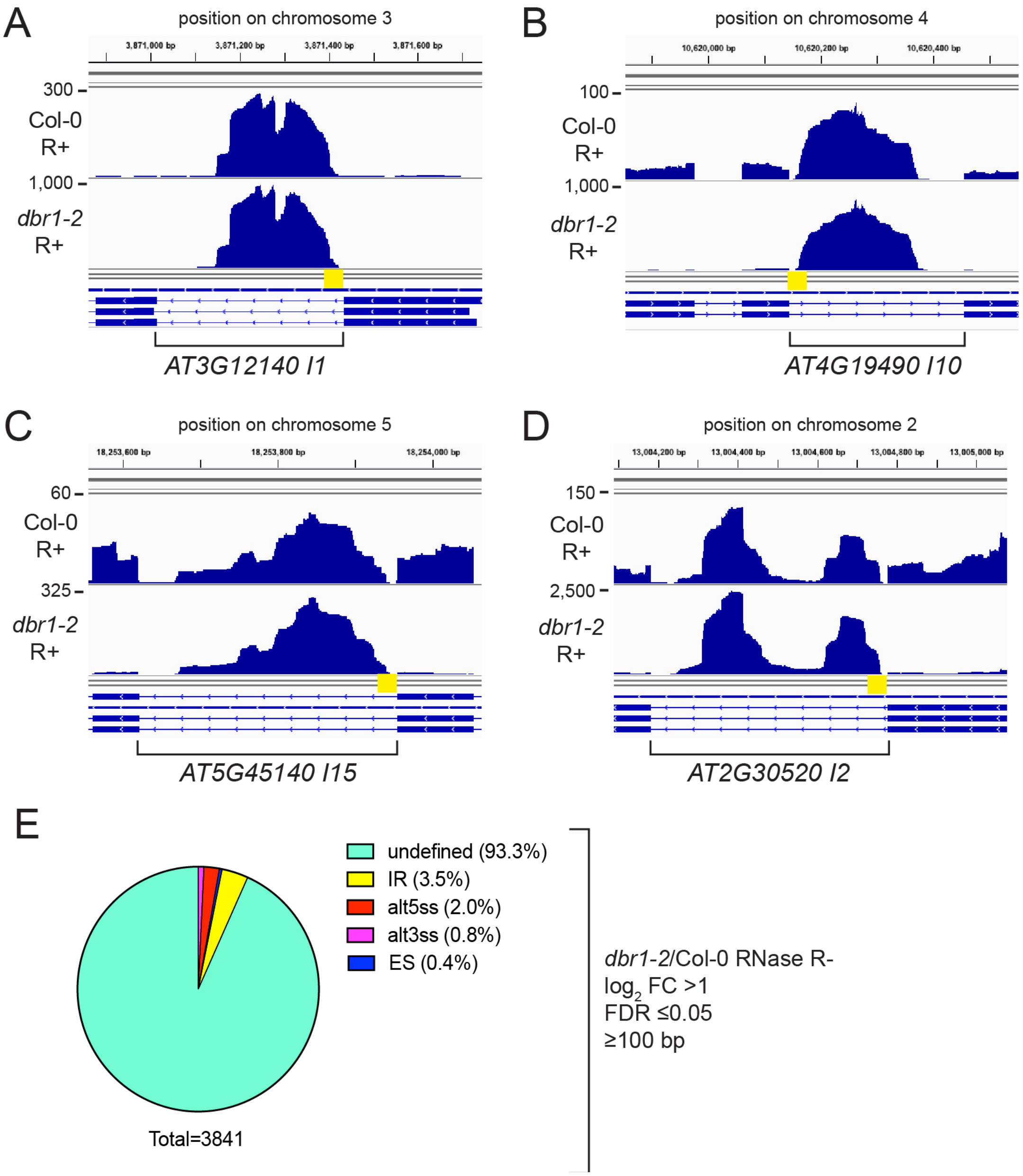


**SFigure 6. ASpli analysis of RNAseq profiling of *dbr1-2* and Col-0.** A-D) Read pileup images for chromosomal positions around *AT3G12140 I1* (A), *AT4G19490 I10* (B), *AT5G45140 I15* (C), and *AT2G30520 I2* (D) from Col-0 (top panel) and *sic-3* (bottom panel) RNA samples treated with RNase R (R+) (Li et al., 2016). Scale is read count and yellow box highlights intron 5'-end. Images were generated by the Integrative Genomics Viewer, version 2.4.1. (Robinson et al., 2011). E) Proportion of all *dbr1-2* differential usage splice variants, relative to Col-0, corresponding to the indicated splice variant types meeting the filtering and statistical criteria shown. The RNA samples for this analysis were not treated with RNase R (RNase R-) (Li et al., 2016).

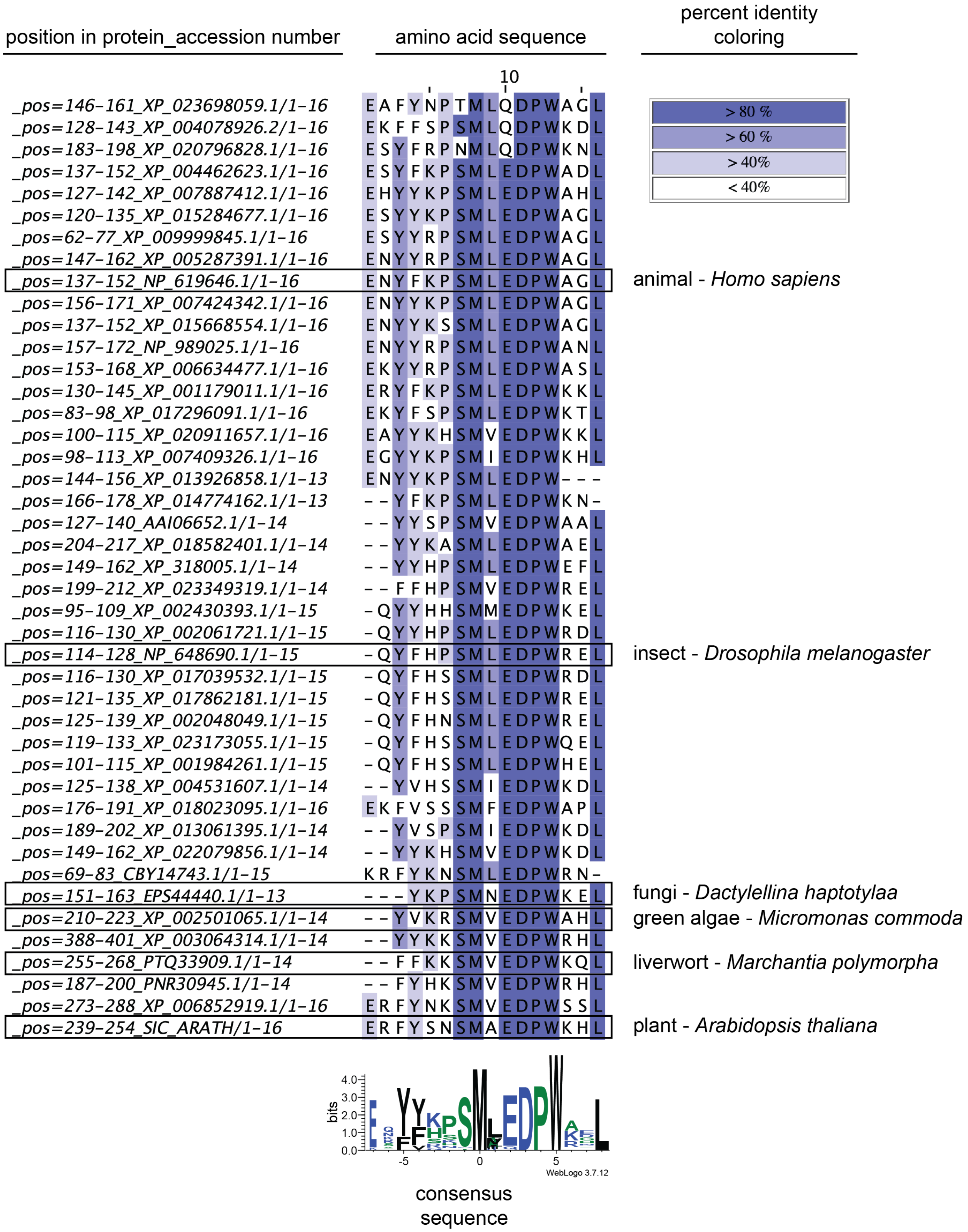


**SFigure 7. Alignment of MPLKIP motifs from plants, fungi, and animals.** The "position in protein_accession number" column indicates the region of the protein corresponding to the

listed National Center for Biotechnology Information Reference Sequence. Amino acid sequences were aligned with MAFFT (Katoh et al., 2005). Sequences are organized according to Kingdom with Animals at top and Plants at bottom, ending with the sequence from *SIC*. Percent amino acid identity is indicated by coloring according to the scale shown. Consensus sequence is shown as a Weblogo (Crooks et al., 2004) with amino acids colored as polar (green), basic (blue), hydrophobic (black), and basic (red).

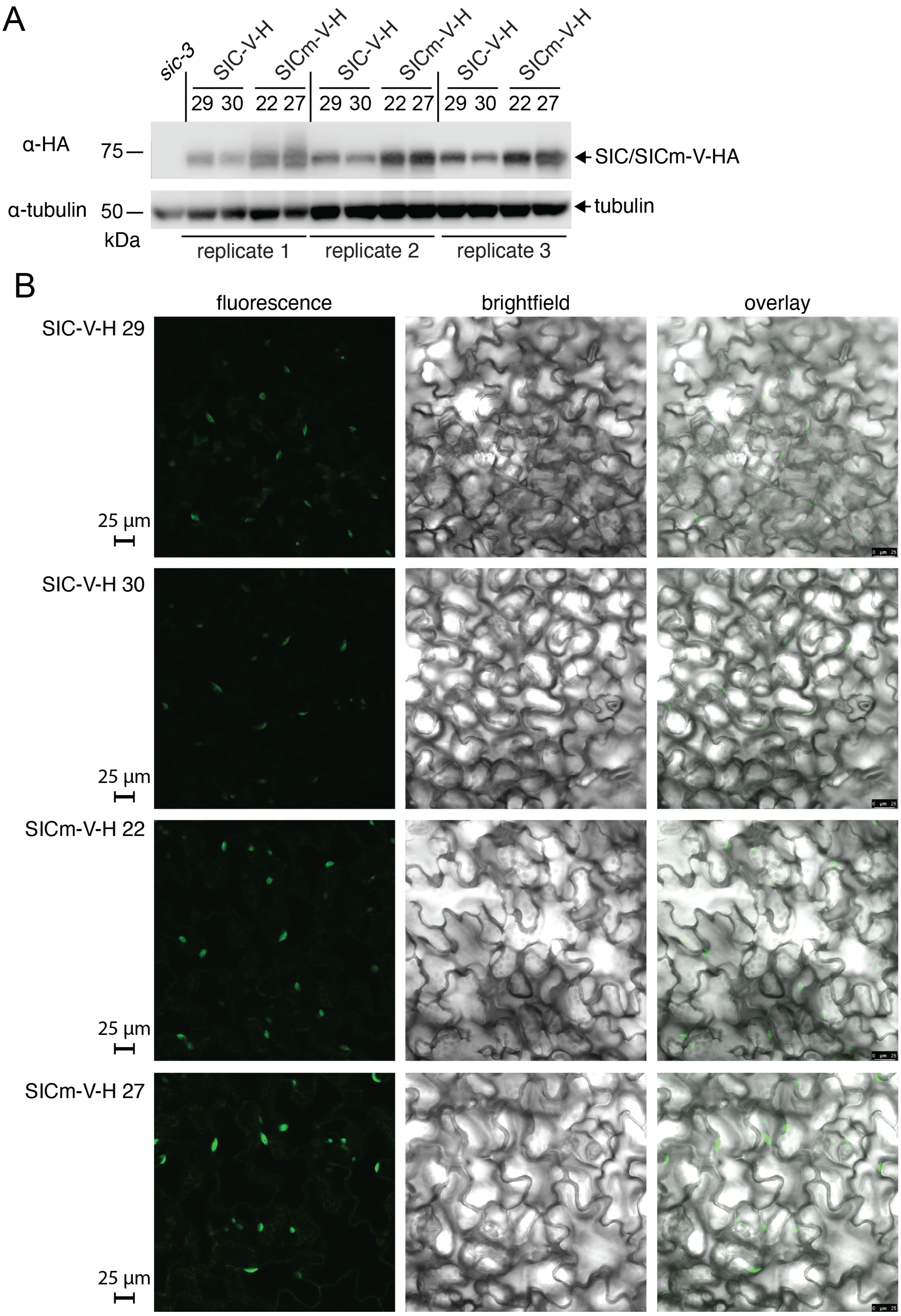
A
sic-3
SIC-V-H
SICm-V-H
SIC-V-H
SICm-V-H
SIC-V-H
SICm-V-H
29 30 22 27 29 30 22 27 29 30 22 27
α-HA
75
SIC/SICm-V-HA
α-tubulin
50
tubulin
kDa
replicate 1
replicate 2
replicate 3
B
fluorescence
brightfield
overlay
SIC-V-H 29
25 µm
SIC-V-H 30
25 µm
SICm-V-H 22
25 µm
SICm-V-H 27
25 µm

**SFigure 8. Characterization of *SICp:SIC-mVenus-HA/sic-3* and *SICp:SICm-mVenus-HA/sic-3* transgenic lines.** A) Western blot analysis of whole cell extracts from *SICp:SIC-mVenus-HA*/*sic-3* and *SICp:SICm-mVenus-HA*/*sic-3* transgenic lines. Top blot shows SIC-mVenus-HA (SIC-V-H) and SICm-mVenus-HA (SICm-V-H) protein from the independent transgenic at the T5 generation. HA-tagged SIC-V-H and SICm-V-H proteins were detected with anti-HA primary antibody and anti-mouse secondary antibody. Bottom blot shows tubulin loading control detected with anti-tubulin primary antibody and anti-mouse secondary antibody. Protein molecular weight standards in kDA are shown at the left of the blot. B) Subcellular localization of SIC-V-H and SICm-V-H proteins in cotyledons of 7-day-old T5 seedlings of the indicated transgenic lines determined by confocal microscopy.t Images labeled "fluorescence" are signal from the YFP channel (emission 520-550 nm) with 514 nm excitation, "brightfield" are from the Transmitted Light Detection channel, and "overlay" are the merge of these two channels. mVenus-tagged fluorescent proteins appear in subcellular structures consistent with nuclei. Scale bars indicate 25 $\mu$m.
